# Prolonged delays in human microbiota transmission after a controlled antibiotic perturbation

**DOI:** 10.1101/2023.09.26.559480

**Authors:** Katherine S. Xue, Sophie Jean Walton, Doran A. Goldman, Maike L. Morrison, Adrian J. Verster, Autumn B. Parrott, Feiqiao Brian Yu, Norma F. Neff, Noah A. Rosenberg, Benjamin D. Ross, Dmitri A. Petrov, Kerwyn Casey Huang, Benjamin H. Good, David A. Relman

## Abstract

Humans constantly encounter new microbes, but few become long-term residents of the adult gut microbiome. Classical theories predict that colonization is determined by the availability of open niches, but it remains unclear whether other ecological barriers limit commensal colonization in natural settings. To disentangle these effects, we used a controlled perturbation with the antibiotic ciprofloxacin to investigate the dynamics of gut microbiome transmission in 22 households of healthy, cohabiting adults. Colonization was rare in three-quarters of antibiotic-taking subjects, whose resident strains rapidly recovered in the week after antibiotics ended. In contrast, the remaining antibiotic-taking subjects exhibited lasting responses, with extensive species losses and transient expansions of potential opportunistic pathogens. These subjects experienced elevated rates of commensal colonization, but only after long delays: many new colonizers underwent sudden, correlated expansions months after the antibiotic perturbation. Furthermore, strains that had previously transmitted between cohabiting partners rarely recolonized after antibiotic disruptions, showing that colonization displays substantial historical contingency. This work demonstrates that there remain substantial ecological barriers to colonization even after major microbiome disruptions, suggesting that dispersal interactions and priority effects limit the pace of community change.

## Introduction

New microbes constantly enter the digestive tract from our diets and environments, but few of these microbes become long-term residents of the adult gut microbiome^1–3^. To colonize the gut, microbes must overcome a series of ecological barriers: they must disperse into a new host, survive the harsh environment of the stomach^4,5^, and compete against resident microbes to occupy an ecological niche^3,6–11^. Understanding these ecological barriers is crucial for understanding how gut bacteria spread between hosts^12–14^ and for the design and persistence of microbiome therapeutics^3,11,15–17^.

Despite these substantial ecological barriers to colonization, people who live together frequently carry pairs of nearly identical microbial strains^12–14,18,19^, suggesting that commensal microbes can transmit between cohabiting partners and colonize established communities. However, few studies have conducted the longitudinal sampling necessary to capture these transmission events in action^13,20^, and it remains unclear what ecological contexts allow microbes to successfully colonize new hosts.

One prominent view is that colonization of the gut is primarily determined by the availability of open niches^21–25^. In mouse models, for example, the presence of a single resident strain can prevent other, closely related species from colonizing, suggesting that niche availability could play a major role in the colonization resistance of the gut microbiota^26–31^. This simple view of colonization predicts that any perturbation that opens niches in an established community should lead to rapid colonization and extensive community change. But in natural human settings, it remains unclear how ecological factors like dispersal limitation^32^ and priority effects^33^ constrain the introduction and initial expansion of new microbes, especially because their effects are difficult to distinguish from a lack of available niches. Disentangling these ecological barriers requires controlled perturbations that open niches in the human gut, along with longitudinal sampling to link these open niches to subsequent colonization dynamics.

To address this gap, we tracked the dynamics of colonization in a household cohort before and after a controlled antibiotic perturbation. Antibiotics are a common perturbation that can disrupt the colonization resistance of established gut communities^23,34–36^. While this property is often used to promote the engraftment of deliberately introduced strains^37,38^, previous work has also shown that gut microbiota can be remarkably resilient to antibiotic perturbations over longer timescales, recovering much of their initial species-level composition in the weeks following treatment^35,39–41^. It remains unclear to what extent this resilience is driven by the recovery of resident strains^41^ or the colonization of new strains, and whether strains shared with cohabiting partners can serve as a reservoir to promote microbiome recovery^42^. We sought to address these questions by using strain-resolved metagenomics to profile the microbiomes of antibiotic-taking subjects and their cohabiting partners at high temporal and genetic resolution. These data allow us to observe the colonization of niches opened by antibiotics, providing a unique window into the ecological forces governing colonization in native human gut communities.

## Results

### Heterogeneous responses to antibiotic perturbation in a household cohort

To measure the dynamics of colonization in a natural human setting, we performed a longitudinal study of 48 healthy adults from 22 households (**Fig. 1a** and **Extended Data Fig. 1**). Subjects collected stool samples at a mixture of weekly and daily intervals for two months, and in the middle of the study, one subject in each household took the broad-spectrum antibiotic ciprofloxacin orally for five days as a controlled perturbation to open niches in the established, adult gut microbiome^23,34,35^. After this initial sampling period, subjects also collected follow-up samples every few months for up to two years after the antibiotic perturbation, allowing us to examine colonization dynamics over longer timescales. We performed metagenomic sequencing of these stool samples and used a reference-based pipeline to profile species abundance and strain-level variation^41,43,44^.

**Fig. 1.**
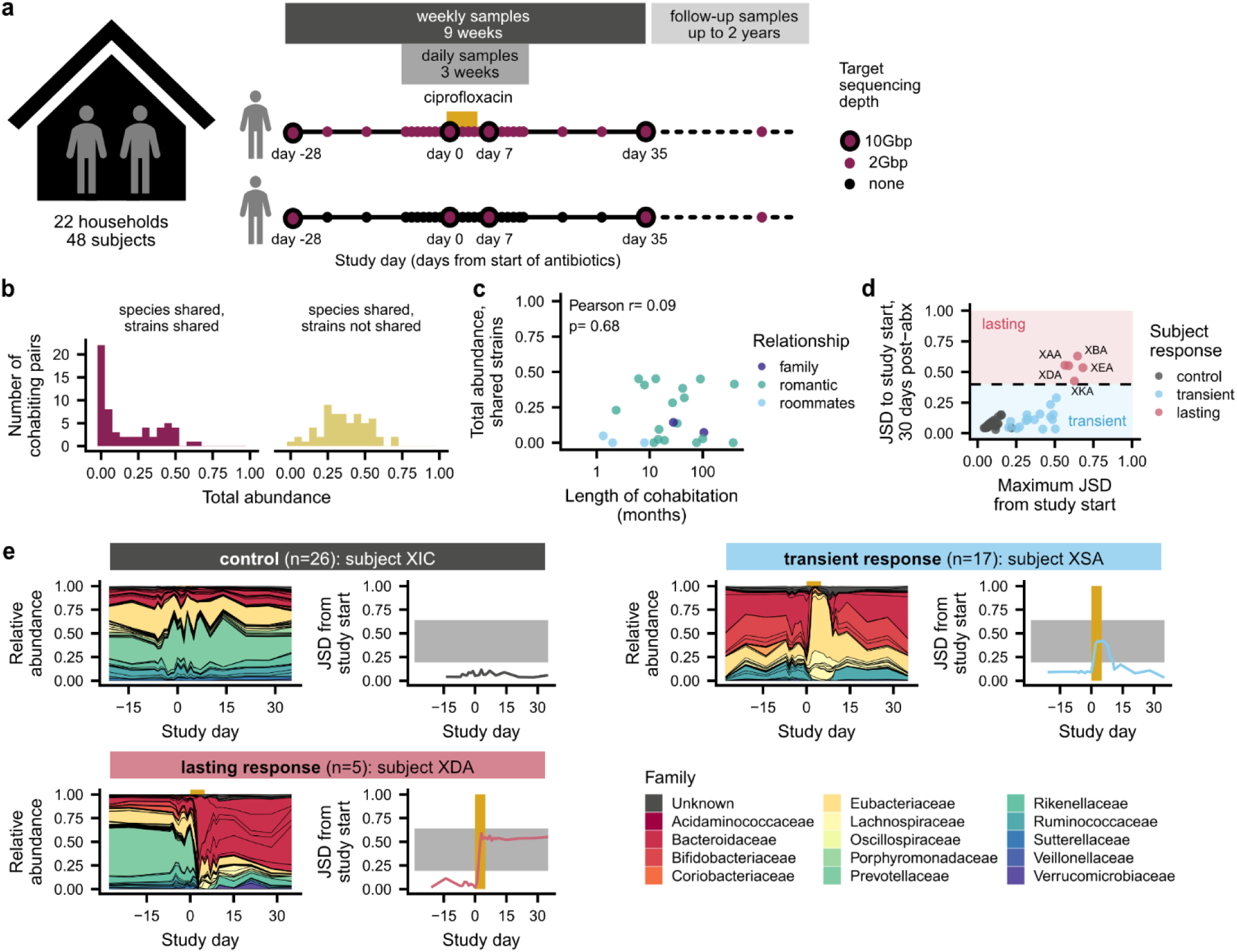
Heterogeneous responses to antibiotic perturbation in a longitudinal household cohort. **a**, Study design. Healthy, cohabiting adults collected weekly stool samples for 9 weeks, during which one member of each household took a 5-day course of ciprofloxacin. In 18 households, subjects collected occasional follow-up samples for up to two years after the initial 9-week sampling period. **b**, Cohabiting subjects shared a range of gut microbial strains. Histograms show the distribution of the total relative abundance of populations with and without shared strains among cohabiting pairs at the initial sampling timepoint. **c**, The total abundance of shared strains was not correlated with the length of cohabitation (Pearson’s test, n=22, ρ=0.09, P=0.68). Points show the longest-cohabiting pair in each household. **d,** Heterogeneous responses to antibiotic perturbation. The antibiotic responses of each community can be classified as transient (blue) or lasting (red) based on the Jensen-Shannon divergence (JSD) of their species composition between the beginning and end of the main study. **e**, Examples of transient and lasting antibiotic responses, relative to a control subject. Left panels show the species-level community composition over time, while right panels show the JSD from the initial timepoint. Gold bars indicate the ciprofloxacin perturbation, while the gray shading illustrates the range of JSDs between non-cohabiting subjects (2.5 to 97.5 percentiles).

Consistent with previous work^12–14,19^, we found that many pairs of cohabiting, unrelated adults shared gut microbial strains at the beginning of the study, indicating that they were acquired through previous horizontal transmission (**Fig. 1b**). To identify shared strains, we first identified species shared between cohabiting partners, and we classified the strains of that species as “shared” if they had higher genetic similarity than 99% of strain pairs of the same species from unrelated subjects (**Extended Data Fig. 2a-e; Methods**; median threshold of >99.9% average nucleotide identity across species, or >75% of private marker SNPs detected). The total amount of strain sharing varied widely across households: in 7 of 30 (23%) cohabiting pairs, we detected no shared strains, whereas in the remaining subjects, we identified a median of four shared strains that comprised a median of 23% of the relative abundance of the gut microbiome (**Fig. 1b** and **Extended Data Fig. 2f,g**). The total abundance of shared strains was not correlated with the length of cohabitation (**Fig. 1c**; Pearson’s test, n=22, ρ=0.09, P=0.68). Two subject pairs shared >30% of their gut microbiomes after living together for less than one year, but we detected no shared strains in two households who had lived together for 6 and 30+ years, respectively (**Fig. 1c**). These data show that shared strains do not accumulate linearly with time, and that even prolonged cohabitation is not sufficient to fully homogenize human gut microbiomes. We hypothesized that these varied amounts of strain sharing could provide a backdrop for investigating the responses to antibiotic perturbation.

**Fig. 2.**
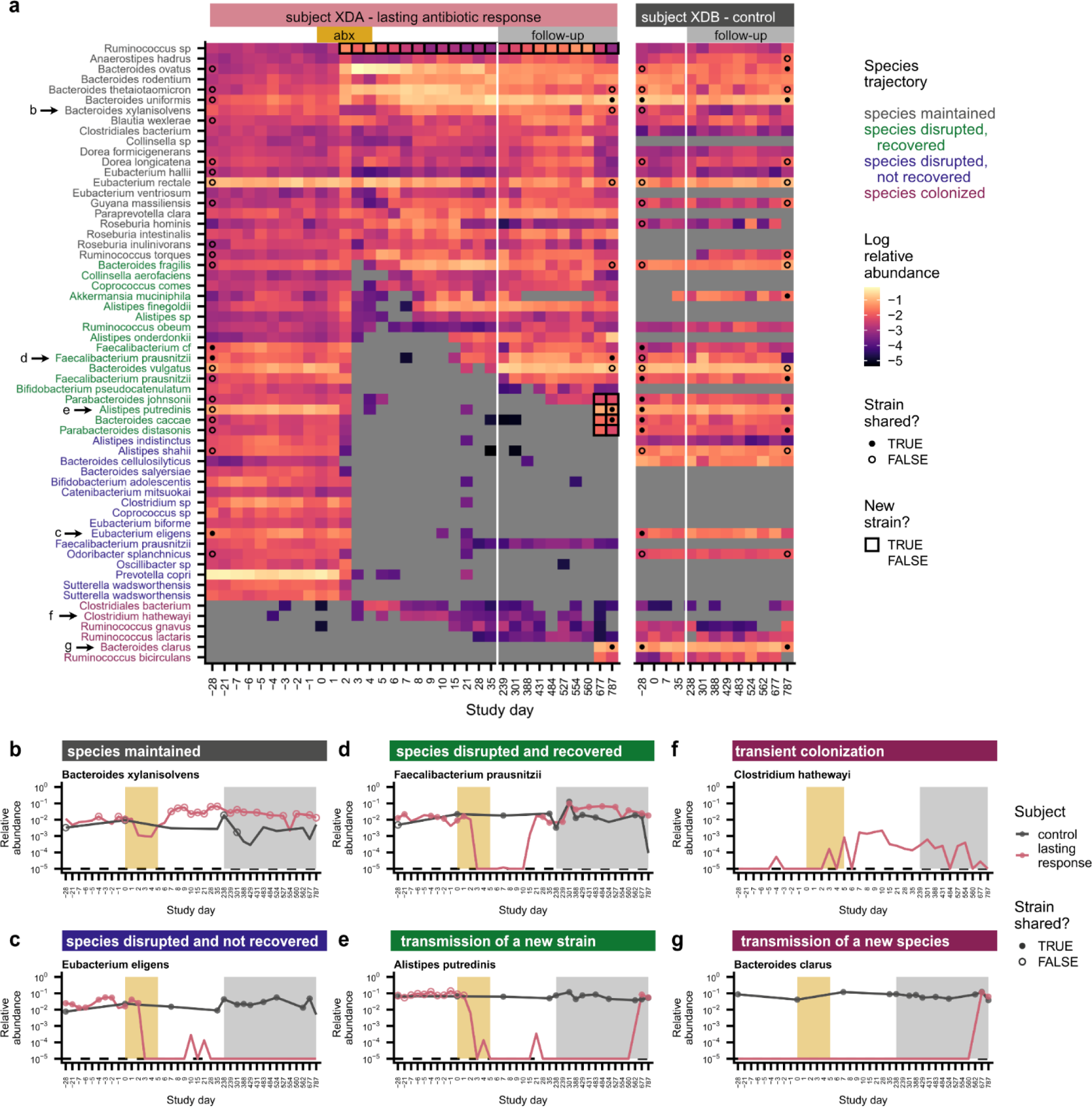
Species disruption, recovery, and colonization in a subject with lasting antibiotic responses. **a**, Relative abundances of species in subject XDA (left) and their cohabiting partner XDB (right). Species are shown if they were present in XDA before antibiotics (median relative abundance >0.1%) or if they newly colonized XDA (**Methods**). Points at the first and last timepoints indicate whether the strains at that timepoint were shared (closed) or not shared (open) with any timepoint from the cohabiting subject (**Methods**); no point is shown if the species was not shared between subjects, or if the sequencing depth was insufficient to determine whether strains were shared. Timepoints are outlined in black if a new strain was detected relative to the initial timepoint. **b-g,** Examples of species from XDA that (**b**) were maintained through ciprofloxacin perturbation; (**c**) were disrupted by ciprofloxacin and did not recover; (**d**) were disrupted by ciprofloxacin and experienced recovery of the resident strain; (**e**) were disrupted by ciprofloxacin and experienced recovery via the transmission of a new strain from the cohabiting partner; (**f**) transiently colonized after ciprofloxacin perturbation; and (**g**) were newly transmitted from the cohabiting partner after ciprofloxacin perturbation. Points indicate whether the strains at that timepoint were shared (closed) or not shared (open) with the strains at any timepoint in the cohabiting partner; points are not shown if the species was not shared between subjects, or if the sequencing depth was insufficient to determine whether strains were shared (**Methods**). The gold and gray boxes denote the ciprofloxacin perturbation and the period of follow-up sampling, respectively. The dashed line indicates the limit of detection, 10^-5^.

We first characterized the community-level effects of ciprofloxacin on gut microbiome composition. Consistent with previous work^45,46^, we found that ciprofloxacin caused rapid changes in community composition that exceeded the magnitude of day-to-day variability in the control subjects, who did not take antibiotics (**Fig. 1d,e**, **Extended Data Fig. 3a**). However, the magnitude and duration of the responses varied widely across subjects. Most antibiotic-taking subjects (n=17/22, 77%) had only transient antibiotic responses and declines in community diversity that lasted 1-2 weeks (**Fig. 1d** and **Extended Data Fig. 3b; Methods**), in line with previous work showing that gut microbiomes often display resilience after antibiotic perturbation^35,39,40^. In contrast, five subjects (n=5/22, 23%) experienced major changes in species composition that persisted for >1 month after ciprofloxacin perturbation (**Fig. 1e** and **Fig. 2**, **Extended Data Figs. 3c,h e4-7**), showing that subjects can have heterogeneous responses to the same antibiotic perturbation.

**Fig. 3.**
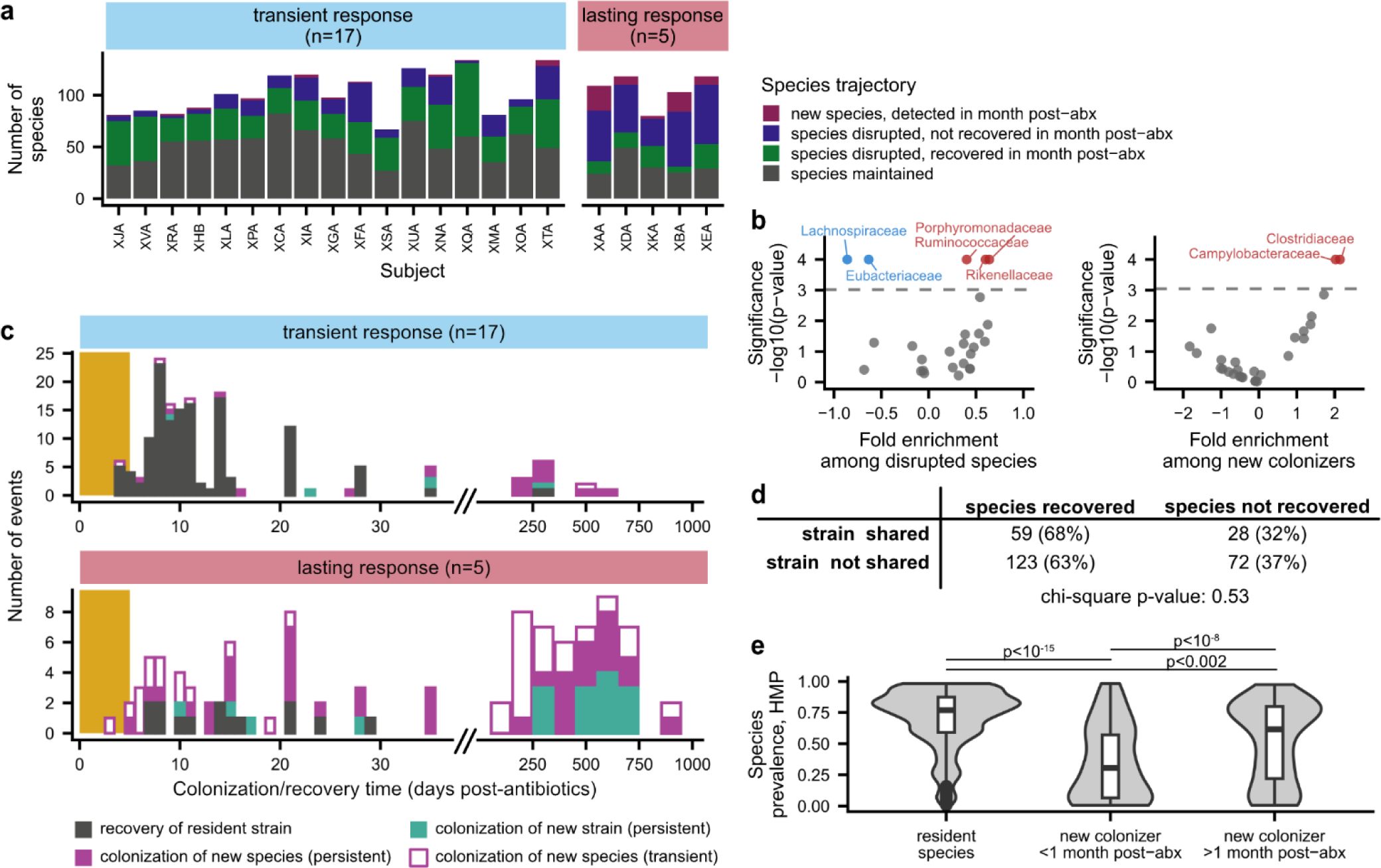
Dynamics of species disruption and colonization in the month after antibiotic perturbation. **a**, Number of species classified as maintained, disrupted, recovered, and colonized after ciprofloxacin in each antibiotic-taking subject. **b**, Enrichment and depletion of bacterial families among disrupted species (left) and new colonizers (right). Families with significant enrichment (red) or depletion (blue) are labeled (permutation test, Bonferroni-corrected P<0.05). **c**, Timeline of species recovery and colonization after antibiotic perturbation. Colonization times were inferred by fitting relative abundance trajectories to a simple model of exponential growth with saturation (**Methods**). Strains and species are annotated as transient colonizers if they no longer had relative abundance >10^-3^ one month after antibiotics. Species that were present before antibiotics but had insufficient coverage after antibiotics to determine strain-level composition are not shown. The gold bar shows the timing of the ciprofloxacin perturbation. **d**, Relationship between strain sharing and species recovery for all antibiotic taking subjects. Strain sharing between cohabiting subjects does not increase the likelihood of species recovery after ciprofloxacin perturbation (chi-square test, n=282, χ^2^=0.4, P=0.53). **e**, New colonizers have lower prevalence in the Human Microbiome Project compared to resident species (Wilcoxon rank-sum two-sided tests, resident species vs. new colonizers <1 month post-antibiotics: n=1333, r=0.22, P=5.0 x 10^-16^; resident species vs. new colonizers >1 month post-antibiotics: n=1376, r=0.16, P=2.7 x 10^-9^; new colonizers <1 month post-antibiotics vs. new colonizers >1 month post-antibiotics: n=161, r=0.26, P=1.2 x 10^-3^). Box plots show the median and interquartile range of each distribution.

**Fig. 4.**
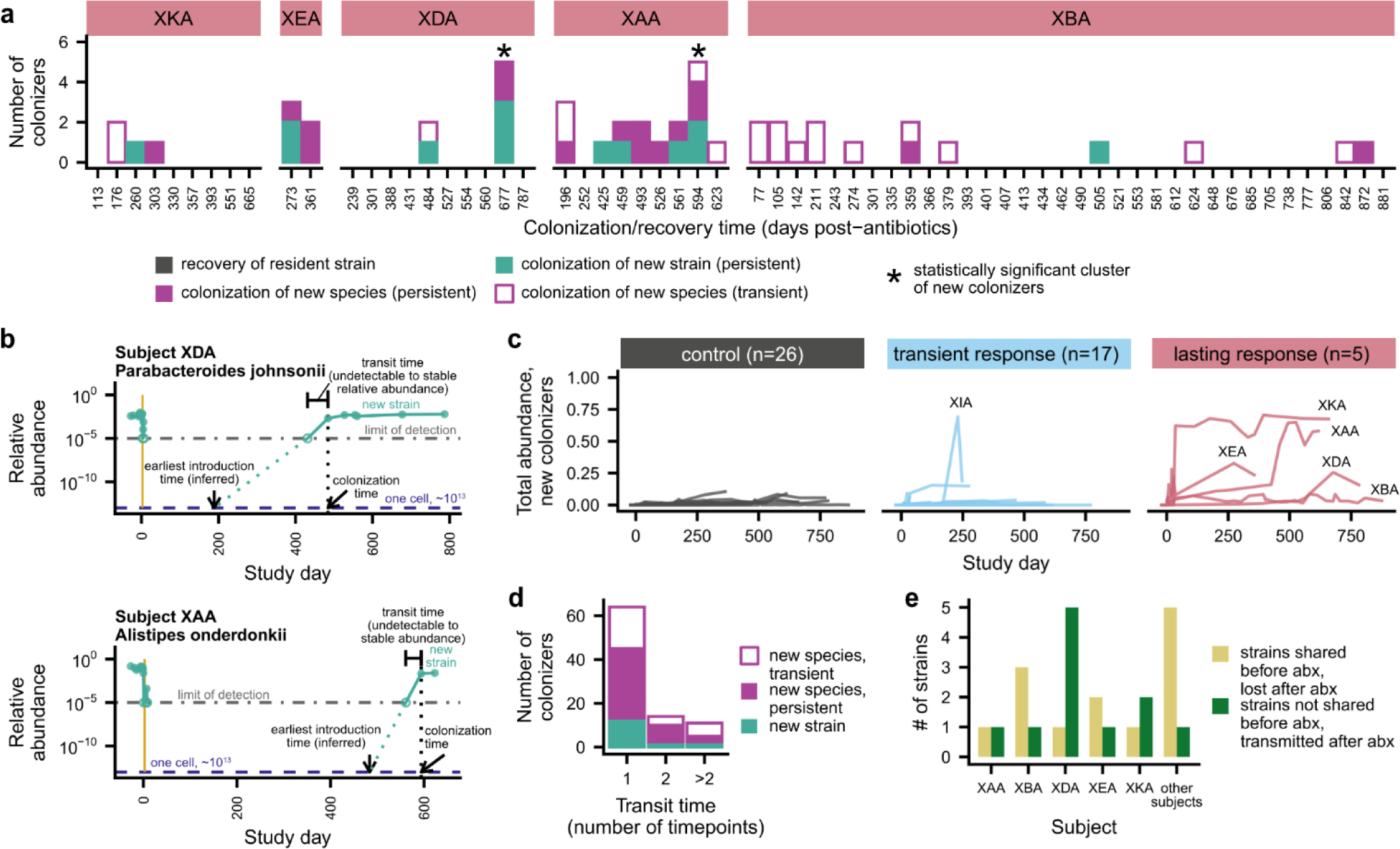
Colonization was elevated in subjects with lasting antibiotic responses, but only after long delays. **a**, Timeline of species recovery and colonization in subjects with lasting antibiotic responses. Colonization times were inferred by fitting relative abundance trajectories to a simple model of exponential growth with saturation (see panel b; **Methods**). Strains and species are annotated as transient colonizers if they no longer had relative abundance >10^-3^ at the end of the study. Timepoints with statistically significant cohorts of colonization events are annotated with an asterisk (permutation test, Bonferroni-corrected P<0.05). Species that were present before antibiotics but had insufficient coverage after antibiotics to determine strain-level composition are not shown. Due to space constraints, timepoints with no new colonization were subsampled to once per month for subject XBA. **b**, The earliest introduction time, transit time, and colonization time of each new colonizer can be inferred from relative abundance trajectories (**Methods**). The gold bar shows the timing of the ciprofloxacin perturbation. These two cases depict the colonization of a new strain of a resident species that was disrupted by antibiotics. **c**, Colonization was elevated in subjects with lasting antibiotic responses in the two years after ciprofloxacin perturbation. Lines show the total relative abundance of newly colonizing strains and species in each subject over time. **d**, New colonizers rapidly reached carrying capacity, even when they colonized long after antibiotics. Histograms show the number of intervals required for each new strain or species to increase from undetectable relative abundances to its inferred carrying capacity. **e**, Strains that are shared with a cohabiting partner can fail to recolonize after antibiotics, even as new strains are acquired from the cohabiting partner. The number of shared strains lost and gained after antibiotic perturbation is shown for each subject with lasting antibiotic responses and in aggregate for all other antibiotic-taking subjects.

We investigated various host and microbial factors that might explain these heterogeneous responses. Subjects with lasting antibiotic responses underwent changes in absolute abundance that were broadly comparable to those in subjects with transient responses, as estimated by colony plating (**Extended Data Fig. 3d; Methods**). We also found no relationship between the antibiotic response and prior antibiotic usage (**Extended Data Fig. 3e**), pre-perturbation community diversity (**Extended Data Fig. 3f**; Wilcoxon rank-sum two-sided test n=22, r=0.28, P=0.22), or the amount of strain sharing at the beginning of the study (**Extended Data Fig. 3g**; Wilcoxon rank-sum two-sided test n=22, r=0.04, P=0.88). Instead, we found that the lasting antibiotic responses were most strongly predicted by the magnitude of the initial perturbation (**Fig. 1d**). This link between perturbation magnitude and duration suggests that sufficiently large perturbations can cross a “tipping point” that overcomes the resilience of the gut microbiome and shifts the community into a new state^39,40,45,46^.

### Rapid recovery of resident strains in subjects with transient antibiotic responses

The heterogeneous responses to ciprofloxacin present an opportunity to link these large-scale changes in community composition to subsequent colonization dynamics. We quantified the species-level dynamics within each community (**Fig. 3**, **Extended Data Fig. 8**), hypothesizing that subjects who experienced more extensive species disruptions would experience elevated levels of colonization and transmission.

Many species experienced large declines in relative abundance (>32-fold) during the 5-day course of ciprofloxacin (**Fig. 2a-e**, **3a**, and **Extended Data Fig. 8a**). In subjects with transient antibiotic responses, these disrupted species accounted for ∼20-80% of the initial gut microbiome composition, compared to <10% in controls (**Extended Data Fig. 8b**). The *Rikenellaceae*, *Porphyromonadaceae*, and *Ruminococcaceae* families were significantly enriched among disrupted species, whereas the *Lachnospiraceae* and *Eubacteriaceae* were significantly less likely to experience disruption (**Fig. 3b** and **Extended Data Fig. 8c**; **Methods**; permutation test, Bonferroni-corrected P<0.05).

In subjects with transient antibiotic responses, a majority of disrupted species recovered to within an order of magnitude of their pre-antibiotic relative abundance (n=574/821, 70%; **Fig. 2d** and **3a**; **Extended Data Fig. 8b**). The genetic similarity of the pre- and post-antibiotic strains indicated that this recovery was driven by the resident strain, or recolonization by a close relative, rather than colonization by an unrelated strain of the same species (**Fig. 3c**). Of the 133 recovered populations with sufficient sequencing coverage to analyze strain-level composition, 129 (97%) involved the recovery of the pre-antibiotic strain, and none of the remaining populations showed evidence of transmission from the cohabiting partner (**Methods**).

Previous studies in mouse models have hypothesized that strain transmission from cohabiting individuals could contribute to these high levels of resident strain recovery^42^. Surprisingly, we found that the rate of recovery was independent of any prior household sharing (**Fig. 3d**; chi-square test for all antibiotic-taking subjects, n=282, χ^2^=0.4, P=0.53; **Extended Data Fig. 8g**, chi-square test for subjects with minimal and transient responses, n=210, χ^2^=0.3, P=0.58). We also observed many cases in which a disrupted species failed to recover more than a month after antibiotics, even when the strain was shared with the cohabiting partner (**Fig. 2c** and **Fig. 3d**; **Extended Data Fig. 8d** and **8e**). These data show that there can be substantial ecological barriers to transmission between cohabiting subjects, even when niches appear to be available in the gut microbiome.

### Resident strain recovery restores colonization resistance in subjects with transient antibiotic responses

We investigated the extent to which these species losses provide opportunities for new microbes to colonize. To identify new colonizers, we searched for newly colonizing species that were not detected at the beginning of the study (**Fig. 2f and 2g**) as well as newly colonizing strains of an existing species that were not detected at the beginning of the study and had high genetic divergence from the initial resident strain (**Fig. 2e**; **Methods**). Despite the many species disruptions that we observed, we detected no new colonizing strains or species during the first month after antibiotics in over half of subjects with transient responses (n=9/17, 53%; **Extended Data Fig. 8h**), and the total relative abundance of newly colonizing strains and species was not significantly different between these subjects and the controls (**Extended Data Fig. 8i**; Wilcoxon rank-sum two-sided test, n=43, r=0.26, P=0.10).

One explanation for the limited colonization in the first month after ciprofloxacin is that it could take new colonizers a longer time to reach detectable relative abundances. We tested this idea by investigating the longer-term dynamics of colonization in follow-up samples collected for up to two years after the antibiotic perturbation. Of the 22 households who completed the main, two-month sampling design, 18 households collected a median of 3 follow-up samples per subject over a median of 12 months after the end of the main study. These data revealed that, even after two years of follow-up sampling, colonization remained limited in subjects with transient antibiotic responses (**Fig. 3c**). The number (median: 0; Wilcoxon rank-sum two-sided test, n=34, r=0.09, P=0.61) and relative abundance (median: 0.4%; Wilcoxon rank-sum two-sided test, n=43, r=0.14, P=0.36) of new colonizers in these subjects remained similar to the control subjects, who also experienced minimal new colonization even over these longer timescales (**Extended Data Fig. 8f, 9a,b**). The limited colonization we observe up to two years after ciprofloxacin perturbation suggests that the rapid recovery of resident strains in subjects with transient antibiotic responses is sufficient to restore the natural colonization resistance of the gut microbiome.

### Extensive species losses and disruptions of colonization resistance in subjects with lasting antibiotic responses

We hypothesized that subjects with lasting antibiotic responses might experience higher levels of new colonization. Although the proportion of disrupted species was only slightly higher in these subjects compared to those with transient responses (**Fig. 3a**, **Extended Data Fig. 8b**), these disrupted species exhibited much lower rates of recovery in subjects with lasting responses: only ∼25% (n=78/309) of disrupted species recovered in the month after the ciprofloxacin course ended, compared to ∼70% (n=574/821) in subjects with transient responses (**Fig. 3a**). Most disrupted species did not recover and instead appeared to be lost from the gut microbiome, even when they were shared with a cohabiting partner (**Fig. 3d**).

Consistent with this hypothesis, many new strains and species reached detectable abundances in the month after antibiotics in subjects with lasting responses (**Fig. 3c** and **Extended Data Fig. 8h**). However, these new species were rarely detected in healthy gut microbiomes: compared to resident species, these new colonizers had significantly lower prevalence (**Fig. 3e**; Wilcoxon rank-sum two-sided test, n=1333, r=0.22, P=5.0 x 10^-16^) and median abundance (**Extended Data Fig. 8j**; Wilcoxon rank-sum two-sided test, n=1333, r=0.19, P=3.1 x 10^-12^) among subjects in the Human Microbiome Project. The *Campylobacteraceae* and *Veillonellaceae* families were enriched among new colonizers (**Fig. 3b**; **Methods**; permutation test, Bonferroni-corrected P<0.05) and included common oral bacteria like *Campylobacter concisus*, *Dialister invisus*, and *Megasphaera micronuciformis*^47–50^ (**Extended Data Figs. 4b** and **5b**). These findings are consistent with prior work showing that oral microbes are frequently transmitted into the gut^51^ and can reach high relative abundances when gut commensals decrease in absolute abundance^52^. We also identified potential opportunistic pathogens^21,53,54^ like *Klebsiella pneumoniae*, *Citrobacter freundii*, and *Streptococcus gallolyticus* (**Extended Data Fig. 5b**), although no subjects reported side effects or illnesses after taking ciprofloxacin. Altogether, many species expansions in the month after antibiotic perturbation were consistent with overgrowth of potential pathogens, a phenomenon that is common after antibiotic perturbations of the gut microbiome and is often ascribed to a disruption of colonization resistance^54^.

**Fig. 5.**
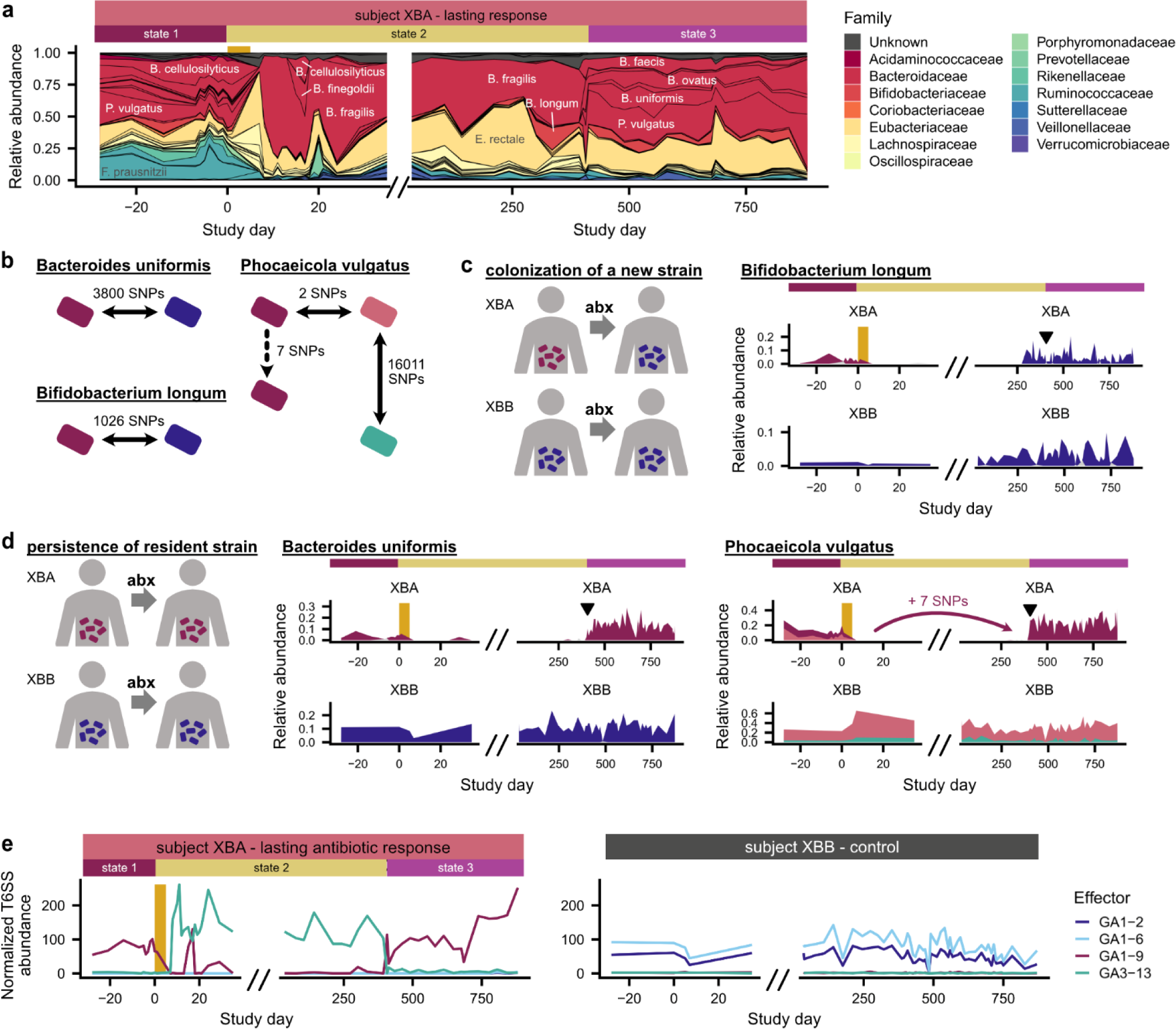
Rapid transitions between alternative states in a subject with a lasting antibiotic response. **a**, Subject XBA underwent rapid shifts in community composition around days 0 and 401. **B**, Number of mutations distinguishing strains of *B. uniformis, P. vulgatus,* and *B. longum* in subject XBA and their cohabiting partner XBB (**Methods**). Strain relative abundances in (**c,d**) were calculated based on the frequencies of these distinguishing SNPs and are shown in corresponding colors. **c**, *Bifidobacterium longum* transmitted from XBB to XBA after antibiotics. **d**, Strains of *Bacteroides uniformis* and *Phocaeicola vulgatus* present before antibiotics re-emerged around day 401. Triangles in (**c,d**) indicate timing of the transition from state 2 to 3. **e**, Type VI secretion system (T6SS) effector genes underwent sharp shifts in abundance that correlate with shifts in community composition.

Although we detected many new species in the month after antibiotics in subjects with lasting antibiotic responses, these new species had limited effects on long-term gut microbiome composition. New species were often transient (**Fig. 2f**), and only 25 of 47 (53%) new species were still detectable a month after the ciprofloxacin course ended (**Fig. 3c**). The primary exception was *Bacteroides stercoris* in subject XKA, which transmitted from the cohabiting partner after ciprofloxacin perturbation and reached a relative abundance of ∼60% by day 64, which it maintained throughout the >1.5 years of follow-up sampling (**Extended Data Fig. 8k**). Overall, however, these examples of commensal colonization were surprisingly rare in the month after antibiotics, suggesting that substantial ecological barriers to colonization can remain even after major microbiome disruptions.

### Commensal strains abruptly colonize subjects with lasting antibiotic responses, but only after long delays

Ecological theory^55,56^ suggests that the low levels of commensal colonization in the subjects with lasting antibiotic responses could be driven by the functional redundancy of the gut microbiome. If the surviving resident species in these subjects can fill the niches cleared by antibiotics, we would expect that colonization would remain limited over longer timescales as well because the gut microbiome has reached a new stable state.

Surprisingly, subjects with lasting responses experienced elevated colonization more than a year after the original perturbation (**Fig. 4a,c** and **Extended Data Fig. 9a,b**; Wilcoxon rank-sum two-sided test, number of colonizers >1 month post-antibiotics, n=26, r=0.67, P=7.0 x 10^-4^). In four of these five subjects, new strains and species comprised ∼15-70% of the gut microbiome one to two years after antibiotic perturbation, compared to only 0.4% in their cohabiting partners (**Extended Data Fig. 9c**; Wilcoxon rank-sum two-sided test, n=11, r=0.77, P=0.014). Of the 25 new colonizers that were still detected at the end of the study and had sufficient sequencing coverage for strain analysis, only one-third (n=8, 32%) were detected in the cohabiting partner, while the rest were acquired from other sources (**Methods**). This elevated colonization in subjects with lasting antibiotic responses suggests that major, antibiotic-mediated disruptions can create niches for new microbes to colonize.

However, while the overall rates of colonization were elevated in these subjects, these colonization events only took place after long delays (**Fig. 4a,b**). We detected 62 colonization events 1-12 months after the perturbation and 50 colonization events 1-2.5 years later (**Fig. 4a**), showing that the timescale of colonization far exceeds the 1–2-week timescale of resident strain recovery (**Fig. 3c**). The large gap between these timescales suggests that new species face substantial ecological barriers to colonization. These long delays are also reminiscent of ecological literature showing that species introduced into new environments often display invasion lags, persisting at low abundances for long periods of time before undergoing exponential expansions^57,58^.

Some delays in the detection of new colonizers are naturally expected due to the time it takes for a species to grow from its initial inoculum to reach detectable relative abundances. Due to the large number of cells in the human gut^59^, these growth lags can be substantial even under strong ecological selection (**Methods**). We can estimate the strength of this effect using our longitudinal data, by comparing the relative growth rates of the invading species with the total time it takes for them to invade (**Fig. 4b, Methods**).

We fit the relative abundance trajectories of new colonizers to a simple ecological model where each species grows exponentially until it reaches its inherent carrying capacity (**Fig. 4b, Methods**). These data revealed that new colonizers expanded rapidly despite their lengthy colonization delays. Of the 112 new strains and species detected more than one month after antibiotic perturbation, 76 (68%) new colonizers increased from undetectable (<10^-5^) to maximum relative abundances between consecutive sampling timepoints (**Fig. 4d**). While the total time between samples varied across our cohort (**Extended Data Fig. 9d**), some of these intervals comprised <5% of the total colonization time and were consistent with a net growth advantage for the colonizer of at least 0.5 doublings per day. We used these inferred relative growth rates to calculate the maximum time required for a newly introduced species to reach carrying capacity, starting from a single cell. Our conservative estimates suggest at least 33 of the 112 (29%) new colonizers were introduced at least six months after the end of antibiotics, with the remaining examples mostly limited by the temporal resolution of our follow-up sampling (**Extended Data Fig. 9d**). This rapid growth shows that new colonizers experience strong ecological selection, and that detection lags cannot fully explain the large colonization delays we observe.

What ecological forces could delay colonization in the face of such strong ecological selection? One hypothesis is that the influx of viable microbes into the gut is extremely low, and the rate of colonization is determined by the rate of these rare microbial introductions. In this case, the inferred introduction times provide a corresponding bound on the overall rates of introduction. For example, a ∼6 month delay after antibiotics (as in **Fig. 4b**) implies an average influx of <0.01 successful migrants per day (**Methods**). However, we also observed strong temporal clustering of colonization events (**Fig. 4a**), suggesting that the simplest models of dispersal limitation do not apply. In subjects XDA and XAA, we identified 5 colonization events in each subject at a single timepoint—more than would be expected to occur by chance if colonization events occurred independently (**Extended Data Fig. 9e**; **Methods**; permutation test, Bonferroni-corrected P<0.05). These clusters of new colonizers often contained groups of closely related species. For example, we simultaneously detected two *Bacteroides* species in XDA (**Fig. 2a**), three *Alistipes* species in XAA (**Extended Data Fig. 4b**), and two new strains of *Phocaeicola vulgatus* in XAA at a different timepoint (**Extended Data Fig. 9f**). This clustering of closely related strains raises the intriguing possibility that facilitative interactions between “guilds” of gut microbial strains^60^ can promote their joint dispersal or colonization and contribute to invasional meltdown^61^, in which invasive species facilitate each other’s establishment.

We wondered whether the strains that were previously shared with the cohabiting partner might play an outsized role in this long-term recovery. However, in 4 of the 5 subjects with lasting antibiotic responses, we found that the recovery of resident strains always occurred within the same narrow time window as in subjects with transient responses (**Fig. 3c**), and, with the exception of subject XBA (see below), we did not identify any instances in which a shared strain recolonized at a later date (**Fig. 4a**). At the same time, we observed several cases in which new strains transmitted from the cohabiting partner after antibiotics, suggesting that the species that failed to recover may also have had opportunities to recolonize (**Fig. 4e**). These findings suggest that ecological factors like priority effects and changing selection pressures limit recolonization, creating historical contingency in microbiome composition.

### Rapid transitions between colonization-resistant alternative states in a subject with a lasting antibiotic response

While four of the five subjects with lasting antibiotic responses experienced elevated levels of new colonization, the remaining subject (XBA) exhibited markedly different dynamics, with new colonizers accounting for only ∼3% of the gut microbiome nearly two and a half years after antibiotics. By sequencing 106 timepoints over a two-year period, we found that XBA’s gut microbiome underwent sudden, striking transitions between three distinct community states (**Fig. 5a** and **Extended Data Fig. 10a**): an initial, diverse state (days -28 to 0), a post-antibiotic state dominated by *Bacteroides fragilis* (days 0 to 401), and a third, diverse state (days 401 to study end). The return to a diverse state, which took place over two weeks, was not associated with any known changes in the subject’s diet, lifestyle, or health status. These long periods of community stability, punctuated by sudden shifts in species abundance, suggest that the gut microbiome can undergo rapid transitions between alternative states^62,63^.

We investigated whether the transition away from *B. fragilis* dominance was driven by the persistence of resident strains or colonization of new strains, including recolonization from the cohabiting partner (**Fig. 5b-d**). One of the first species to return after antibiotics was *Bifidobacterium longum*, which became detectable again around day 301 (**Fig. 5a**). The post-antibiotic strain of *B. longum* was more genetically similar to the strain in the cohabiting partner than the pre-antibiotic strain in XBA, indicating that its expansion was driven by colonization rather than strain persistence (**Fig. 5b,c** and **Extended Data Fig. 10c; Methods**). *B. longum* can inhibit *B. fragilis* in co-culture^64^ and reduce its microbial load *in vivo*^65^, suggesting that colonization by *B. longum* may have contributed to the end of *B. fragilis* monodominance.

After *B. longum* colonized, eight *Bacteroidaceae* species underwent coordinated expansions between days 393 and 407, driving the return to the diverse state. In contrast to *B. longum*, the strains of *B. uniformis* and *P. vulgatus* that we detected after antibiotics were more similar to the pre-antibiotic strains in XBA than strains in the cohabiting partner (**Fig. 5d** and **Extended Data Fig. 10d-f**), indicating that these strains persisted at low abundance through *B. fragilis* monodominance before undergoing rapid, simultaneous expansions. These dynamics bear striking similarities to the delayed, rapid, clustered colonization dynamics that we observed in the other four subjects with lasting antibiotic responses, suggesting that they may be driven by similar ecological forces.

Curiously, we found that the changes in community composition in XBA coincided with sharp shifts in the relative abundance of Type VI secretion system (T6SS) genes (**Fig. 5e** and **Extended Data Fig. 10g-i**). Many *Bacteroidaceae* carry T6SS gene cassettes, which contain effector and immunity genes that allow cells to kill neighboring cells that lack the immunity gene^66–69^ and may contribute to the maintenance of distinct community states^66^. Intriguingly, although XBA and XBB shared many *Bacteroidaceae* strains, their communities had different dominant E-I pairs (**Fig. 5e**), and we speculate that this incompatibility may have limited the ability of *Bacteroidaceae* to transmit and colonize XBA after ciprofloxacin perturbation. From this observational study alone, we cannot determine whether the changing abundances of T6SS genes are the cause or consequence of the rapid switches between alternative community states, but these analyses support the theory that interbacterial antagonism contributes to community stability and colonization resistance in the gut microbiome^53,70^.

## Discussion

Our results show that strong ecological barriers limit colonization in the adult gut microbiome, even after antibiotic perturbations that cause extensive species losses. These findings counter the prevailing view that colonization is primarily limited by the availability of open niches, suggesting instead that the introduction and initial expansion of new microbes also present major barriers to colonization in natural settings.

What ecological forces are responsible for these prolonged delays in colonization after antibiotic perturbation? The rapid colonization dynamics we observe (**Fig. 4b**), together with the elevated levels of colonization in subjects with lasting antibiotic responses (**Fig. 4c**), suggest that new colonizers experience strong ecological selection. A common explanation for colonization delays is that dispersal limitation strongly restricts the pool of available colonizers, slowing the pace of colonization^7^. However, the coordinated clusters of colonization events that we observe are unlikely to arise from random introductions of individual species (**Fig. 4a** and **Extended Data Fig. 9e**), evidence against the simplest models of dispersal limitation.

We argue instead that collective effects shape colonization dynamics by causing microbial fitness to depend strongly on aspects of community context like population size and interactions with other microbes. These collective effects could arise during dispersal—for example, if microbial species migrate together in larger propagules, a form of community coalescence^71^ (**Fig. 4a** and **Extended Data Fig. 9e**). They can also arise after dispersal, creating priority effects^33,72^ in the resident community. Priority effects can occur through inhibitory interactions like interbacterial antagonism, which allows established strains to kill new arrivals^53,73^ (**Fig. 5e**), or through facilitative interactions like quorum sensing^74^, cross-feeding^75^, and the production of public goods^76^. Regardless of their mechanism, strong collective effects in the gut microbiome complicate the design of microbiome therapeutics by creating historical contingency in colonization outcomes (**Fig. 4e**). Large inoculation doses and tailored microbial consortia may be necessary to overcome these ecological barriers and successfully colonize established microbial communities.

This work shows how the combination of controlled perturbations, longitudinal sampling, and strain-resolved metagenomic sequencing can disentangle the ecological forces that shape colonization in natural settings. Our conclusions are limited by the observational nature of our study, as well as the difficulty of detecting colonization events at low relative abundances.

Despite these limitations, *in situ* studies like ours are necessary for investigating how colonization is shaped by uniquely human aspects of lifestyle and gastrointestinal anatomy, factors that are difficult to replicate in mouse models^42,77^ or *in vitro* communities^78^. Future work that combines our approaches with barcoded strain libraries^79^ or spatially resolved sampling devices^80^ can further illuminate the ecological and evolutionary forces that shape colonization in the human gut.

## Materials and methods

### Data availability

Sequencing data will be made publicly available from the NCBI SRA upon manuscript publication.

### Code availability

The computer code that performs the analysis is available at https://github.com/ksxue/household-transmission-after-abx.

### Study design

This study was approved by the Stanford University Institutional Review Board as Protocol 54715, and written, informed consent was obtained from all participants. Healthy, cohabiting adults were recruited from Stanford University and the surrounding community (**Extended Data Fig. 1a-d**). Exclusion criteria included antibiotic usage in the past three months, past reactions to fluoroquinolone antibiotics, chronic illness, pregnancy or nursing, and age under 18. During the main, two-month sampling period, participants collected weekly stool samples for 9 weeks (days -28, -21, -14, -7, 0, 7, 14, 21, 28, 35) and daily stool samples for 19 days in the middle of the study (days -7 to 11) (**Extended Data Fig. 1e**). One participant in each household took a 500mg oral dose of ciprofloxacin twice daily for 5 days (days 0-4). At each time point, participants collected one stool swab that was stored in 1mL RNAlater (Thermo Fisher AM7021) and collected approximately 20mL of raw stool in a sterile vial. Stool samples were frozen immediately after collection in subjects’ home freezers and were stored there for up to four weeks until they were transferred to storage at -80 degrees C in the laboratory. A total of 48 participants in 22 households completed the sampling protocol. Participants from 18 households collected follow-up samples using the same methods at occasional intervals for up to two years after the main sampling period (**Extended Data Fig. 1e**).

Participants also completed a questionnaire at the beginning of the study that included questions about demographics, diet, lifestyle, recent travel and antibiotic use, household structure, and cohabitation history. At the end of the study and every three months during follow-up sampling, participants were asked to report changes in diet, lifestyle, and living arrangements, as well as recent travel, illnesses, injuries, and antibiotic usage.

### DNA extraction and metagenomic sequencing

DNA was extracted from stool swabs stored in RNAlater using the DNeasy PowerSoil HTP 96 Kit (Qiagen 12955-4). To avoid cross-contamination that could lead to erroneous inferences of strain sharing^81^, samples collected from cohabiting partners during the main, two-month sampling period were arrayed on different 96-well plates. When processing follow-up samples, which were fewer in number, samples were arrayed so that no sample was adjacent to a sample from a cohabiting partner. For stool swabs that yielded less than 1.5ng/uL of DNA, we aliquoted a stool fragment from the accompanying sample of frozen, raw stool and redid the DNA extraction using the DNeasy PowerSoil Kit (Qiagen 12888-100).

Libraries were prepared for metagenomic sequencing using the Nextera DNA Flex Library Prep kit (Illumina 20018705). Samples collected at four “key” timepoints (days -28, 0, 7, and 35) from both control and antibiotic-taking subjects were sequenced to a target depth of 10Gbp. All remaining samples collected from antibiotic-taking subjects during the main, two-month sampling period were sequenced to a target depth of 2Gbp. All follow-up samples collected from both control and antibiotic-taking subjects were sequenced to a target depth of 2Gbp. In total, we sequenced 942 samples collected from the 48 subjects in our study (**Extended Data Fig. 1e**).

### CFU counts

We drilled cores from stool samples frozen at -80 degrees C using the CXT 353 Benchtop Frozen Aliquotter (Basque Engineering) and aliquoted 50-100mg of frozen stool from each sample. We resuspended each stool fragment in 0.5mL reduced PBS and plated 10-fold dilutions on Brain-Heart Infusion agar plates supplemented with 5% sheep’s blood in an anaerobic chamber (Coy Instruments). We incubated plates at 37 degrees C and counted colonies after 48 hours.

### Species and SNP profiling of metagenomic samples

We first used skewer^82^ to trim sequencing adapters and bowtie2^83^ to filter out reads that mapped to the human genome. Remaining reads will be made available in the SRA upon manuscript publication. To obtain species abundances, we used the MIDAS^43^ species module to map sequencing reads against a database of universal, single-copy bacterial genes. We used these sequencing reads to calculate species abundances, which were used to compute Jensen-Shannon divergence and normalized FST^84^, a metric of compositional variability.

Using these species abundance estimates, we generated a list of species that were detected in a given subject across all sequenced timepoints. We then used the MIDAS snps module to map sequencing reads against a set of dereplicated reference genomes representing the detected species. For each population, defined by the set of reads in a sample that mapped to a species’ reference genome, we calculated the relative abundance of single-nucleotide polymorphisms (SNPs) at every genome site. To limit the influence of reads with high similarity to multiple reference genomes, we discarded sites that had sequencing coverage more than two-fold above or below the median genome coverage.

### Identifying cross-contamination and other sample abnormalities

To identify cross-contamination during DNA extraction and library preparation that could confound our identification of shared strains between subjects, we calculated the number of fixed differences between each pair of populations with a median sequencing coverage >5, and we identified all unusually similar population pairs, defined as population pairs from non-cohabiting subjects that had fewer than 3 fixed differences. We identified samples that had unusually similar population pairs for multiple species, suggesting that this population similarity was likely to be due to cross-contamination rather than genetic similarity between strains.

In an initial analysis, we identified 24 samples that had signatures of cross-contamination. Most of these samples were located along the edge of a single DNA extraction plate, suggesting that improper sealing of this plate during DNA extraction may have contributed to cross-contamination^81^. The remaining samples with signatures of cross-contamination were in adjacent wells on either their DNA extraction or metagenomic library preparation plates. We removed these samples from downstream analyses.

We re-sequenced these samples by aliquoting a stool fragment from the banked sample of frozen, raw stool, extracting DNA using the DNeasy PowerSoil Kit (Qiagen 12888-100), and preparing libraries for metagenomic sequencing using the Nextera DNA Flex Library Prep kit (Illumina 20018705). We re-calculated the number of fixed differences between each pair of populations with median sequencing coverage >5 after including these re-sequenced samples in our dataset. We identified no samples from non-cohabiting subjects that had unusually similar population pairs for multiple species, suggesting that the level of cross-contamination between our remaining samples was lower than our limit of detection.

We also identified one case in which we suspected that subjects swapped identifiers. Household XH reported that subject XHC took ciprofloxacin. However, XHC had fewer species disruptions than any other antibiotic-taking subject, and we detected more disrupted species in XHB than in XHC (**Extended Data Fig. 8a** and **8b**). We therefore classified subject XHB as the antibiotic-taking subject from this household. The main conclusions of our study are not affected by whether we designate XHB or XHC as the antibiotic-taking subject.

### Classifying subject responses to antibiotic perturbation

For each subject, we calculated the Jensen-Shannon divergence (JSD) between the species abundances in the initial sampling timepoint and each subsequent sequenced timepoint. We calculated the maximum JSD from the initial timepoint between days -28 and 35, as well as the final JSD from the initial timepoint to day 35. Antibiotic-taking subjects were classified as having a transient antibiotic response if they had a final JSD <0.4 and a lasting antibiotic response if they had final JSD >0.4.

### Identifying shared strains

We used two complementary methods to infer whether a pair of populations shared strains. In the first, “fixed differences” method, which we adapted from InStrain^85^, we calculated the number of fixed differences between each pair of populations with median sequencing coverage >5. We define a fixed difference as a site at which the two populations carried distinct major alleles (frequency>0.5), and the major allele in each population was detected in the other population at a frequency below the Illumina sequencing error rate of 10^-3^. We determined that two populations shared strains if there were fewer fixed differences between them than in 99% of population pairs from non-cohabiting subjects, and we classified two populations as not sharing strains if the number of fixed differences between them exceeded this threshold (**Extended Data Fig. 2a,b**). We classified strain sharing as unknown if neither population in a pair had median sequencing coverage >5.

We also developed a second, “strain fishing” method for identifying shared strains using “private marker” SNPs^41,44^. We first identified “quasi-phaseable” (QP) populations in which we could infer the genotype of the dominant strain^44^. We classified a population as QP if it had median sequencing coverage >5 and we detected SNPs with frequency between 0.2 and 0.8 at <0.1% of the genome, meaning that the population had low genetic variation. In each QP population, we identified private marker sites at which the major allele in the QP population was the minor allele (frequency<0.5) in all populations of the same species analyzed by Garud, Good *et al*.^44^, which includes the Human Microbiome Project and several additional cross-sectional cohorts.

We determined that two populations shared strains if either population was QP and we detected the QP allele at any frequency above the Illumina sequencing error rate of 10^-3^ at >75% of private marker sites in the other population. We classified two populations as not sharing strains if the QP allele was detected above a frequency of 10^-3^ at <75% of private marker sites (**Extended Data Fig. 2c,d**). We classified strain sharing as unknown if neither population in a pair was QP.

We found that the fixed differences and strain fishing methods agreed on >90% of population pairs that could be analyzed using both methods (**Extended Data Fig. 2e**), and we combined the calls from both methods to generate our final classifications of strain sharing. Strain sharing in a population pair was classified as “unknown” if neither method was able to classify strain sharing or if the two methods returned discordant results; “not shared” if one or both methods classified the population pair as not sharing strains; and “shared” if one or both methods classified the population pair as sharing strains.

### Species disruption, recovery, and colonization

We classified species responses to antibiotics using their relative abundance dynamics. A species was classified as “disrupted” if its median pre-antibiotic relative abundance exceeded 10^-3^ and its minimum post-antibiotic abundance declined by more than 1.5 orders of magnitude below its median pre-antibiotic relative abundance. A disrupted species was also classified as “recovered” if its median post-antibiotic relative abundance in a three-sample window exceeded one-tenth of its median pre-antibiotic relative abundance. We calculated the time of recovery to be the first timepoint in this window during which the species exceeded one-tenth of its median pre-antibiotic relative abundance. Species whose median pre-antibiotic relative abundance exceeded 10^-3^ and were not classified as “disrupted” were classified as “maintained.”

A species was classified as “colonizing” if it was not detected at the first three sequenced timepoints and its median relative abundance in a three-sample window subsequently exceeded 10^-3^. We calculated the time of detection to be the first timepoint in this window during which the species exceeded 10^-3^ relative abundance.

### Colonization of new strains of an existing species

We also sought to identify cases in which new strains of existing species colonized the gut microbiome. We used the methods described above to determine whether strains were shared between a population at the initial and subsequent timepoints from each subject (“**Identifying shared strains**”). Populations were classified as candidate colonizing strains when all timepoints past a certain timepoint (designated as the time of strain colonization) were classified as strains “not shared” or “unknown” when compared with the initial timepoint, suggesting that the initial strain was no longer detected.

We reverified candidate colonizing strains through longitudinal strain analysis of candidate new colonization cases. As before, we calculated the relative abundance of single-nucleotide polymorphisms (SNPs) at every genome site for each population. To limit the influence of reads with high similarity to multiple reference genomes, we discarded sites that had sequencing coverage more than 2.5-fold above or below the median genome coverage. We only kept sites that passed this criterion for at least 80% of samples measured for that population. We found SNPs that underwent selective sweeps by identifying SNPs that grew from a frequency below 0.2 to frequency above 0.8 between a pair of timepoints. We excluded sweeping SNPs where the coverage ratio of the site to the median depth of the genome changed by more than a factor of two between the two timepoints. A strain turnover event occurred if at least 1000 SNPs swept between a pair of timepoints, indicating that a minor strain rose from low to high frequency.

We searched each population for strain turnover events in which a low-frequency strain that was not initially present swept to high frequency. We calculated the frequency of the minor strain as the median frequency of the SNPs associated with the sweeping strain. We accepted the new colonization call if and only if the frequency of the minor strain was zero at the initial timepoint.

### Mathematical modeling of colonization delays

To interpret the colonization and recovery dynamics we observed, we fit the relative abundance trajectory of each species to a simple mathematical model, in which the resident population grows exponentially at a constant rate 𝑟 before reaching an intrinsic carrying capacity 𝐾. We allowed the growth rates and carrying capacities to vary between species, and between different populations of the same species in different hosts. Each trajectory in this model can therefore be specified by a triplet of parameters, 𝑟, 𝐾, and 𝜏, where 𝜏 is the time that the population first reaches carrying capacity (the “colonization time” in **Fig. 3c** and **4a**).

We estimated the model parameters by minimizing the sum of squared errors between the observed and predicted values of the logarithm of the relative abundance. We only included timepoints where the observed relative abundance of the species exceeded our limit of detection, which we estimated as the inverse of the total marker gene coverage summed across all species in that sample. We also restricted the model fit to the time interval where the first colonization (or re-colonization) event was estimated to have occurred. To eliminate potential degeneracies in the inferred parameters, we added a weak regularization term to the loss function,

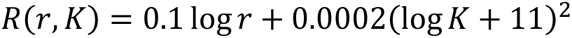

which favors the smallest values of 𝑟 and 𝐾 that are consistent with the observed data. Error minimization was performed in a custom Python script using the minimize function from the SciPy library^86^.

Using these inferred parameters, we estimated the total time required for each species to grow to its final carrying capacity starting from a single founding cell. Under simple models of demographic noise^87^, this time can be approximated by the asymptotic formula,

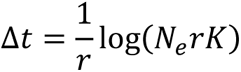

where 𝑁_𝑒_ is the effective population size of a single person’s gut microbiome. To be conservative, we used an effective population size of 𝑁_𝑒_∼10^13^, which corresponds to the total number of bacterial cells in a typical person’s gut^59^. This large effective population size implies that the total growth times can be quite long. For example, a growth advantage of 𝑟 = 1 per day and a carrying capacity of 𝐾∼0.01 corresponds to a total growth time of Δ𝑡∼25 days.

If the estimated value of Δ𝑡 is less than 𝜏, then the difference 𝜏 − Δ𝑡 represents the earliest possible time that the colonizing cell was introduced. Under the assumption that successful colonizers arrive as a Poisson process, the expected value of 𝜏 − Δ𝑡 is equal to the inverse of the rate of successful introductions. We used this relation to convert the minimum possible introduction time into a corresponding bound on the rate of introduction.

### Statistical tests of taxonomic enrichment

We used permutations to test whether certain bacterial families were enriched among disrupted species (**Fig. 3b**). To test for taxonomic enrichment among disrupted species, we began with a simple null model in which species from all bacterial families are equally likely to be disrupted by ciprofloxacin. To generate a null distribution of the number of disrupted species in each family, we identified all populations with median pre-antibiotic relative abundance above 10^-3^ in the ciprofloxacin-taking subjects, and we drew populations at random without replacement, matching the total number of disrupted populations observed across all ciprofloxacin-taking subjects. We calculated the number of disrupted populations in each bacterial family in this simulated dataset, and we compared this distribution to the number of disrupted populations in each bacterial family observed in our ciprofloxacin-taking subjects. To calculate the p-values shown in the text, we calculated the fraction of simulations with a more extreme number of disrupted populations in each family than the observed value and performed a Bonferroni correction by dividing by the number of families tested.

We used an analogous approach to test for taxonomic enrichment among colonizing species (**Fig. 3b**) and species that were disrupted by antibiotics but failed recover (**Extended Data Fig. 8d**).

### Comparing the prevalence and abundance of resident species and new colonizers in the Human Microbiome Project

We downloaded the species abundance profiles analyzed in Good and Rosenfeld^88^, which were generated by using the MIDAS^43^ species module to map sequencing reads from the Human Microbiome Project^89^ (HMP) against a database of universal, single-copy bacterial genes. We calculated the prevalence of each species, defined as the number of subjects in which the species was detected at any non-zero frequency, as well as the median relative abundance of each species in subjects that carried the species at a non-zero frequency.

We identified new colonizers as described above. We also identified a set of resident species, which were detected at any non-zero frequency at the beginning of the study and subsequently attained a median relative abundance >10^-3^ in a three-sample window. These resident species fulfilled the same abundance criteria that we used to identify new species, but, in contrast to new species, they were detected at the beginning of the study. We identified the HMP prevalence (**Fig. 3e**) and median relative abundance (**Extended Data Fig. 8j**) of each resident species and new colonizer in each subject.

### Inferring co-colonization of P. vulgatus strains in subject XAA

We observed that the pre- and post-antibiotic *P. vulgatus* populations in XAA harbored thousands of SNPs at intermediate frequencies, suggesting co-colonization by two or more strains in both the pre- and post-antibiotic intervals. Our strain-sharing analysis above demonstrated that the major strains turned over between the pre- and post-antibiotic timepoints, so we sought to determine whether the minor strains were new colonizers as well.

To carry out this analysis, we assigned individual SNPs to 4 strain haplotypes by manually clustering their allele frequency trajectories over time. We employed the same filters on SNPs and timepoints that we used for verifying the pre-existing strain calls above. We also polarized the SNPs such that the allele frequency referred to the minor allele in the cohort used for calling private marker SNPs above. A manual examination of the allele frequency spectrum at the initial timepoint suggested that the resident population was composed of two diverged strains in a 60:40 ratio. We assigned SNPs to the major strain (strain 1) if their frequency in the initial timepoint fell between 60% and 95%, and to the minor strain (strain 2) if their frequency was between 5% and 40%.

We assigned SNPs to the post-antibiotic strains based on their frequencies in two key timepoints. Day 459 was a quasi-phaseable (QP) timepoint, and was thus well-suited for identifying the major strain. We assigned SNPs to the major strain in the post-antibiotic period (strain 3) if their frequency was >80% this timepoint. Similarly, we found that the sample from one day in the post-antibiotic period (day 425) had a large number of SNPs between 30-70%, and was thus well-suited for identifying candidate SNPs that might belong to the minor strain. We therefore assigned SNPs to the minor strain in the post-antibiotic time window (strain 4) if their frequency at day 425 was >20%, but their frequency in all subsequent timepoints was <50%. We estimated the total number of genetic differences between strains (**Extended Data Fig. 9f**) by taking the differences between the lists of SNPs assigned to each strain.

We estimated the relative frequencies of the strains by taking the characteristic frequencies of the SNPs assigned to each cluster. Since a fraction of these SNPs will be shared across multiple strains, we restricted these estimates to the private marker SNPs in each cluster, which were inferred using the same procedure as above (“**Identifying shared strains**”). To mitigate the effects of mapping errors, we only considered SNPs in the list of core genomes, as previously described^44^. We also excluded SNPs that were present in both the major and minor strains in the post-antibiotic period. This procedure yielded a total of 32, 7, 17, and 23 private SNPs for strains 1-4, respectively. We estimated the frequencies of the respective strains as the median frequency of the private SNPs corresponding to that strain. The median frequency of the pre-antibiotic strains was zero in the post-antibiotic timepoints, and vice versa, confirming that the re-emergence of the *P. vulgatus* population was caused by the colonization of two new strains.

### Distinguishing between strain re-emergence and strain replacement in subject XBA

We sought to determine whether increases in the relative abundance of *Bifidobacterium longum* and several *Bacteroidaceae* species in subject XBA after antibiotics was caused by the reemergence of pre-antibiotic strains or recolonization from XBA’s cohabiting partner, XBB. We first identified species that had at least one QP population (median sequencing coverage >5, SNPs with frequency between 0.2 and 0.8 at <0.1% of the genome; see **Identifying shared strains**) before antibiotics and after antibiotics in both XBA and XBB. Of the 13 species shown in **Extended Data Fig. 10a**, *Bacteroides uniformis, Phocaeicola vulgatus, Bacteroides faecis,* and *Bifidobacterium longum* met these criteria.

#### Bacteroides uniformis

We identified 3800 sites in the core genome of *B. uniformis* at which the major allele in XBA (frequency>0.8) was the minor allele in XBB (frequency<0.2) in all pre-antibiotic, QP populations. At these sites, the major allele in XBA before antibiotics, which we called the XBA allele, had median frequency 1 in XBA and median frequency 0 in XBB before antibiotics. We called the strain carrying the XBA allele at these sites “strain 1,” and we called the strain carrying the alternate allele “strain 2.” The XBA allele was present at a median frequency of 1 at these sites after antibiotics in XBA and 0 in XBB, suggesting that the increase in relative abundance of *B. uniformis* was driven by the re-emergence of strain 1 rather than recolonization by strain 2 (**Fig. 5d** and **Extended Data Fig. 9d**). We identified zero sites in the core genome at which the major allele in XBA before antibiotics (frequency>0.8) became the minor allele after antibiotics (frequency<0.2), suggesting that there were no SNPs that reached fixation between the end of antibiotics at day 5 and the re-emergence of *B. uniformis* around day 401.

#### Phocaeicola vulgatus

We identified two sites in the core genome of *P. vulgatus* at which the major allele in XBA (frequency>0.8) was the minor allele in XBB (frequency<0.2) in all pre-antibiotic, QP populations. At these two sites, the major allele in XBA before antibiotics, which we called the XBA allele, had median frequency ∼0.85 in XBA and median frequency 0 in XBB before antibiotics. We called the strain carrying the XBA allele at these two sites “strain 1,” and we called the strain carrying the alternate alleles “strain 2.” The XBA allele was present at a median frequency of 1 after antibiotics in XBA and 0 in XBB, suggesting that the increase in the relative abundance of *P. vulgatus* was driven by the re-emergence of strain 1 (**Fig. 5d** and **Extended Data Fig. 9e**).

We also identified and manually verified 7 sites in the core genome at which the major allele in XBA before antibiotics (frequency>0.8) became the minor allele after antibiotics (frequency<0.2). At these 7 sites, the major allele in XBA after antibiotics had a median frequency of 1 in XBA and 0 in XBB, suggesting that these SNPs reached fixation on the background of strain 1 between the end of antibiotics at day 5 and the re-emergence of *P. vulgatus* around day 401 (**Fig. 5d** and **Extended Data Fig. 9f**). These sites were widely dispersed across the genome, suggesting that they represent distinct mutations rather than a single recombination event.

Based on the site-frequency spectrum of the *P. vulgatus*, we inferred that the XBB population consisted of at least two distinct, co-colonizing strains. We identified 16011 sites in the genome that had intermediate allele frequencies (0.2<frequency<0.8) in all non-QP populations from XBB and had correlated frequencies across samples, suggesting that these sites distinguish the two co-colonizing strains. Earlier, we determined that the majority strain in XBB differed from the majority strain in XBA at only two segregating sites. We designated the majority strain in XBB, which carried the XBA-like alleles at these 16011 sites, “strain 2,” and we called the minor strain in XBB “strain 3.” To calculate the frequency of strain 3, we calculated the median frequency of SNPs at these 16011 sites (**Fig. 5d**).

#### Bifidobacterium longum

We identified 3667 sites in the core genome of *Bifidobacterium longum* at which the major allele in XBA (frequency>0.8) was the minor allele in XBB (frequency<0.2) in all pre-antibiotic, QP populations. At 1026 of these sites, the major allele in XBB before antibiotics, which we called the XBB allele, had median frequency 0 in XBA and 1 in XBB before antibiotics, suggesting that these sites distinguish the dominant strains in each subject. (We suspect that the remaining sites distinguish a minor, co-colonizing strain, but our sequencing data were insufficient to estimate the frequency of this strain.) We called the strain carrying the XBA allele at these 1026 sites “strain 1,” and we called the strain carrying the XBB allele at these sites “strain 2.” The XBB allele was present at a frequency of 1 after antibiotics in XBA and 1 in XBB, suggesting that colonization of XBA by strain 2 was responsible for the increase in the relative abundance of *B. longum* after antibiotics (**Fig. 5c** and **Extended Data Fig. 9c**).

#### Bacteroides faecis

We identified 9 sites in the core genome of *Bacteroides faecis* at which the major allele in XBA (frequency>0.8) was the minor allele in XBB (frequency<0.2) in all pre-antibiotic, QP populations. These 9 sites were widely dispersed across the genome, suggesting that they represent distinct mutations rather than a single recombination event. One group of four SNPs had correlated frequency trajectories, suggesting that they were carried on a single haplotype (SNP group 1). The remaining five SNPs also had correlated frequency trajectories, suggesting that they were carried on a second, distinct haplotype (SNP group 2). Based on the SNP frequency trajectories and the pigeonhole principle, we inferred that the XBA and XBB populations were comprised of three distinct strains: strain 1, which carried the initial XBA major allele at all 9 sites; strain 2, which carried the initial XBB major allele at all 9 sites; and strain 3, which carried the initial XBA major allele at the four sites in SNP group 1 and the initial XBB major allele at the five sites in SNP group 2. XBA was co-colonized by strains 1 and 3 before antibiotics, but strain 3 appears to have reached fixation after antibiotics. Strain 3 was also present at a median frequency of ∼0.1 in the *B. faecis* population of XBB throughout the sampling period, so we were unable to determine whether the increase in abundance of *B. faecis* after antibiotics was driven by the re-emergence of strain 3 in XBA or colonization from XBB. Aside from the sites in SNP group 2, we identified zero sites in the core genome at which the major allele in XBA before antibiotics (frequency>0.8) became the minor allele after antibiotics (frequency<0.2), suggesting that there were no SNPs that reached fixation between the end of antibiotics at day 5 and the re-emergence of *B. faecis* around day 401.

### Dynamics of Type VI Secretion Systems

Metagenomic sequencing reads were mapped using bowtie2^83^ against a database of 104 Type VI effector, immunity, and structural genes encompassing three distinct T6SS subtypes from *Bacteroidales* species^66^. To prevent cross-mapping, we included only the final 400 bases of the C-terminal domains of the effector genes. To calculate the normalized abundance of each T6SS gene, we divided the number of reads mapping to the gene by the length of the gene and the total number of reads in the sample and multiplied this number by 10^9^.

## Acknowledgements

The authors thank Susan Mello and Robert Pesich for their assistance with study coordination and Ania Robaczewska for technical assistance with library preparation. We also thank Emily Ebel, Jaime Lopez, Saria McKeithen-Mead, Matthew Olm, Tadashi Fukami, and members of the Petrov, Huang, Good, and Relman labs for valuable discussions and comments on the manuscript. Sequencing support for this project was provided as part of the Chan Zuckerberg Biohub Microbiome Initiative, and additional samples were sequenced at the DNA Services Lab, Roy J. Carver Biotechnology Center, University of Illinois at Urbana-Champaign. K.S.X. was supported by a James McDonnell Foundation Postdoctoral Fellowship in Understanding Dynamic and Multi-Scale Systems and a Jane Coffin Childs Memorial Fund Postdoctoral Fellowship. S.J.W. was supported by a Hertz Foundation Fellowship and a Stanford Graduate Fellowship. The authors acknowledge funding from NIH/NIGMS R35-GM142685 (to B.D.R.); NIH/NIGMS R35-GM118165 (to D.A.P.); NSF EF-2125383 (to K.C.H.); Alfred P. Sloan Foundation grant FG-2021-15708 (to B.H.G.) and NIH/NIGMS R35-GM146949 (to B.H.G.); the Thomas C. and Joan M. Merigan Endowment at Stanford University (to D.A.R.); NIH/NIAID R21-AI168860 (to D.A.R.); and NIH/NIAID R01-AI147023 (to D.A.R.). D.A.P., K.C.H., and B.H.G. are Chan Zuckerberg Biohub Investigators.

## Author contributions

K.S.X, D.A.G., D.A.P., K.C.H., B.H.G., and D.A.R. designed the research; K.S.X., D.A.G, A.B.P., F.B.Y., and N.F.N. performed the research; K.S.X., S.J.W., D.A.G., M.L.M., A.J.V., A.B.P., N.A.R., B.D.R, and B.H.G. analyzed the data; K.S.X., D.A.P., K.C.H., B.H.G., and D.A.R. wrote the paper; and all authors reviewed it before submission.

## Extended Data Figures

**Extended Data Figure 1.**
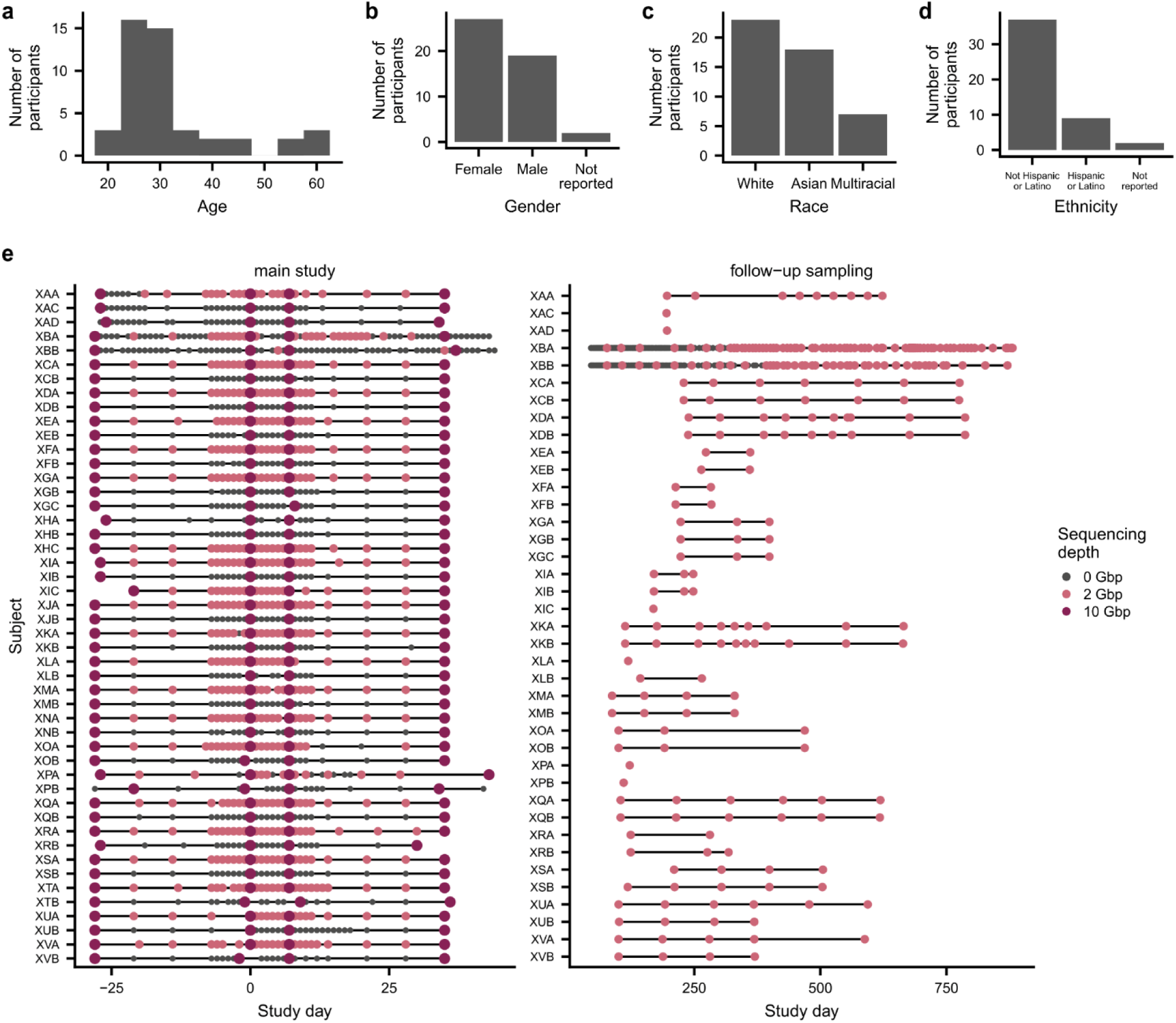
Subject demographics and sampling timelines. Subject demographics, including **a**, age at the beginning of the study; and self-reported **b**, gender; **c**, race; and **d**, ethnicity according to US Census Bureau categories. **e**, Sampling timelines show each sample collected during the study and the target depth to which each sample was sequenced. The second letter of each subject identifier represents the subject’s household; for instance, subjects XAA, XAC, and XAD lived in household A.

**Extended Data Figure 2.**
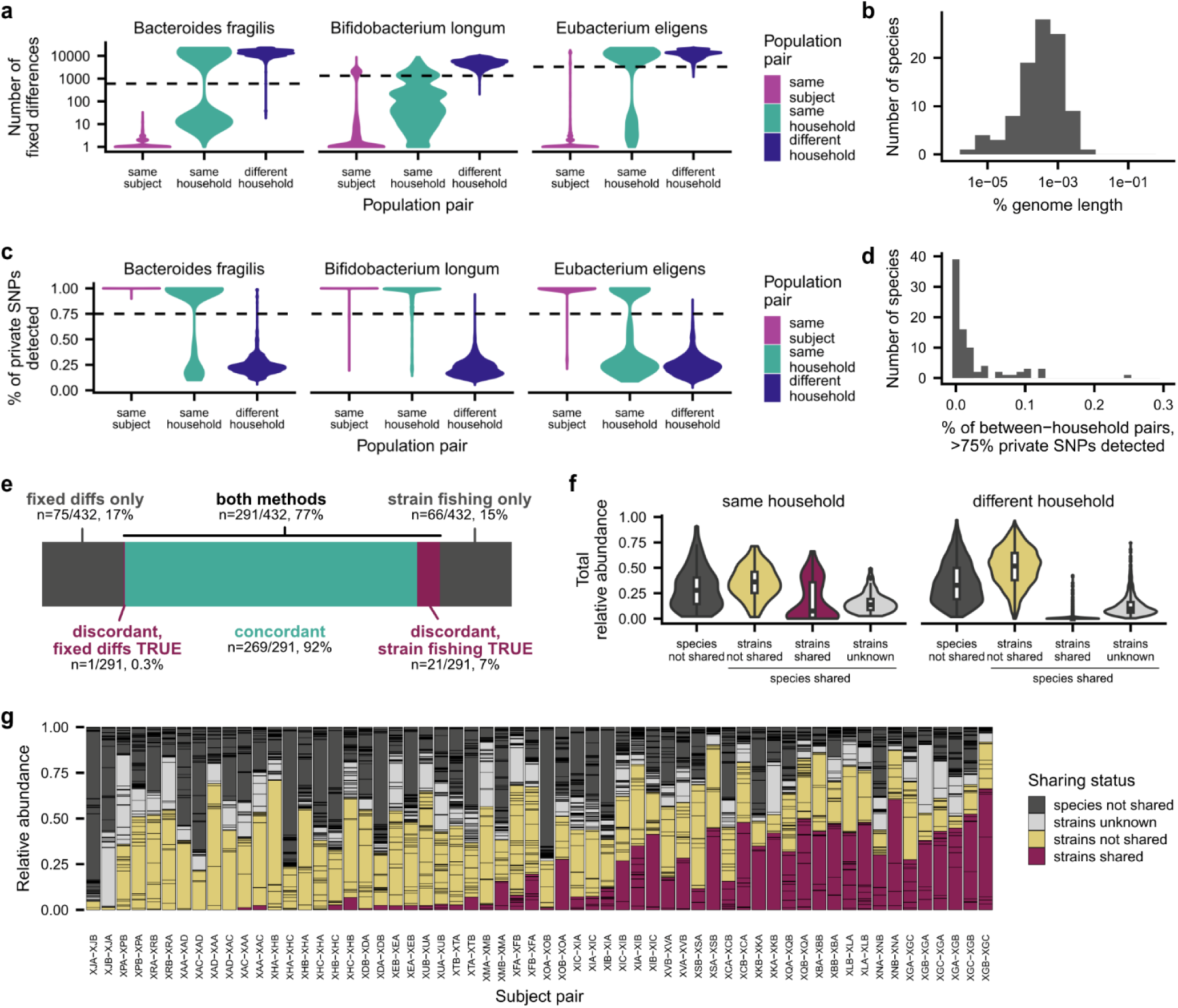
Identifying strains shared between cohabiting subjects. **a**, Number of fixed differences between population pairs for three representative species. Using the “fixed differences” method, we classified two populations from cohabiting subjects as sharing strains if they had fewer fixed differences than 99% of population pairs from non-cohabiting subjects, as shown by the dashed line (**Methods**). Population pairs from the same subject are shown as a control. **b**, Distribution of genetic distances used as a threshold to annotate strain sharing for each species using the “fixed differences” method. **c**, Percent of private SNPs detected in population pairs for three representative species. Using this “strain fishing” method, we classified two populations as sharing strains if we detected >75% of private marker SNPs from one population in the other population, as shown by the dashed line (**Methods**). **d**, Percent of non-cohabiting population pairs for each species that shared >75% of private marker SNPs. **e**, The fixed differences and strain fishing methods agreed on >90% of population pairs that could be analyzed using both methods. **f**, Total abundance of species and strain sharing in pairs of cohabiting and non-cohabiting subjects at the beginning of the study. **g**, Gut microbiome composition at the beginning of the study, with each species colored based on strain and species sharing between cohabiting subjects. In subject pair XLA-XLB, for example, the gut microbiome of subject XLA is shown and colored based on species and strain sharing with subject XLB.

**Extended Data Figure 3.**
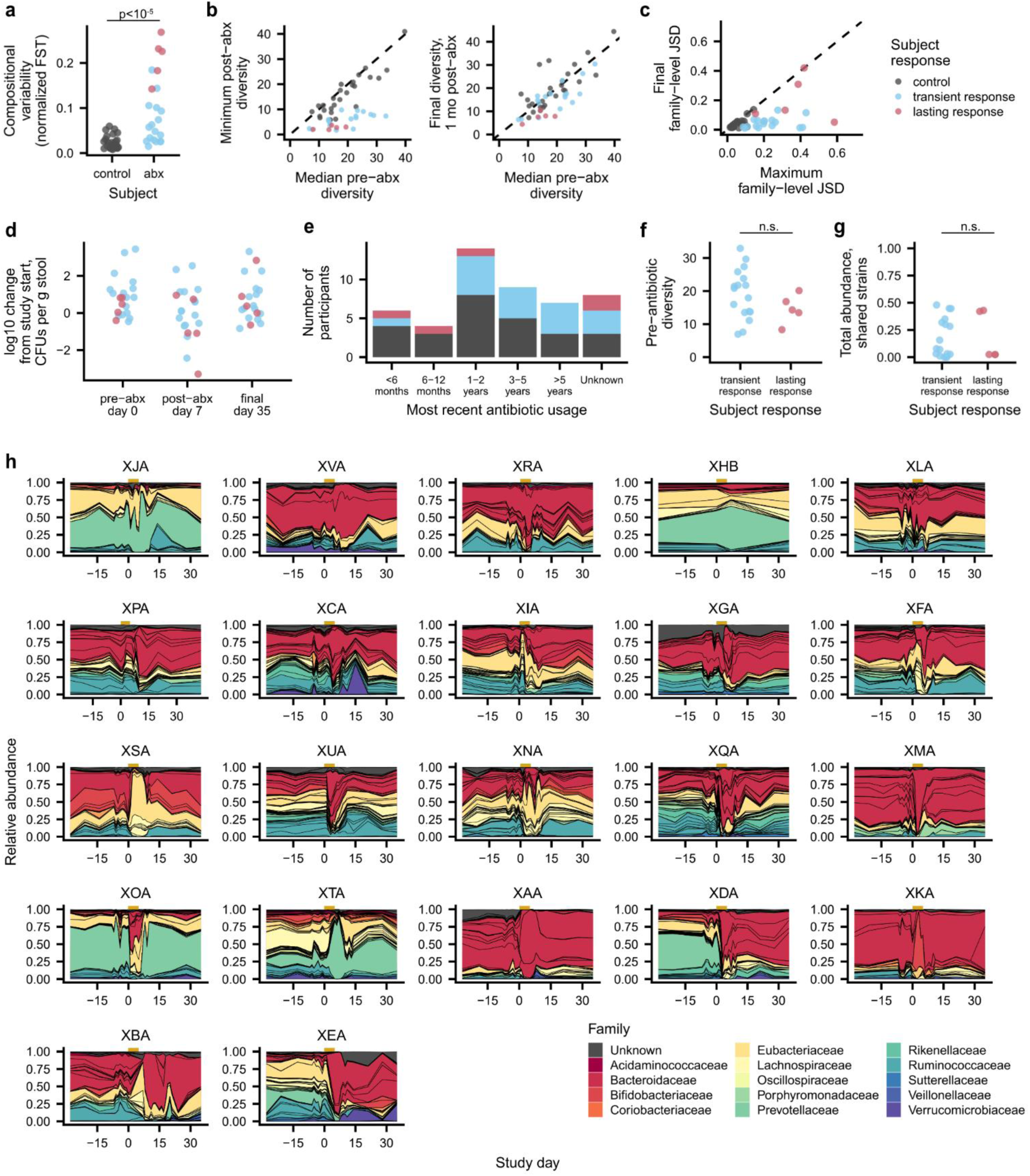
Community responses to antibiotic perturbation. **a**, Antibiotic perturbation increased the compositional variability of the microbiome. Points show the compositional variability of each community calculated at days -28, 0, 7, and 35 using normalized FST^84^ (Wilcoxon rank-sum two-sided test, n=48, r=0.63, P=3.3 x 10^-6^). **b**, Antibiotic perturbation caused mostly transient declines in community diversity. Points show the effective number of species calculated from the Shannon diversity index before antibiotics, after antibiotics, and one month post-antibiotics. **c**, Maximum and final family-level Jensen-Shannon divergence of each community from the initial timepoint during the main study. **d**, Change in CFUs from the initial timepoint. **e**, Most recent antibiotic usage (self-reported) for each subject, colored by subject responses to ciprofloxacin perturbation. **f**, Subjects with transient and lasting responses had similar levels of pre-antibiotic diversity, as quantified by the effective number of species calculated from the Shannon diversity index (Wilcoxon rank-sum two-sided test, n=22, r=0.28, P=0.22). **g**, Subjects with transient and lasting responses had similar proportions of the gut microbiome that were comprised of shared strains (Wilcoxon rank-sum two-sided test, n=22, r=0.04, P=0.88). **h**, Species-level community composition over time for each antibiotic-taking subject. Subjects are arranged in order of increasing maximum Jensen-Shannon divergence from the initial timepoint.

**Extended Data Figure 4.**
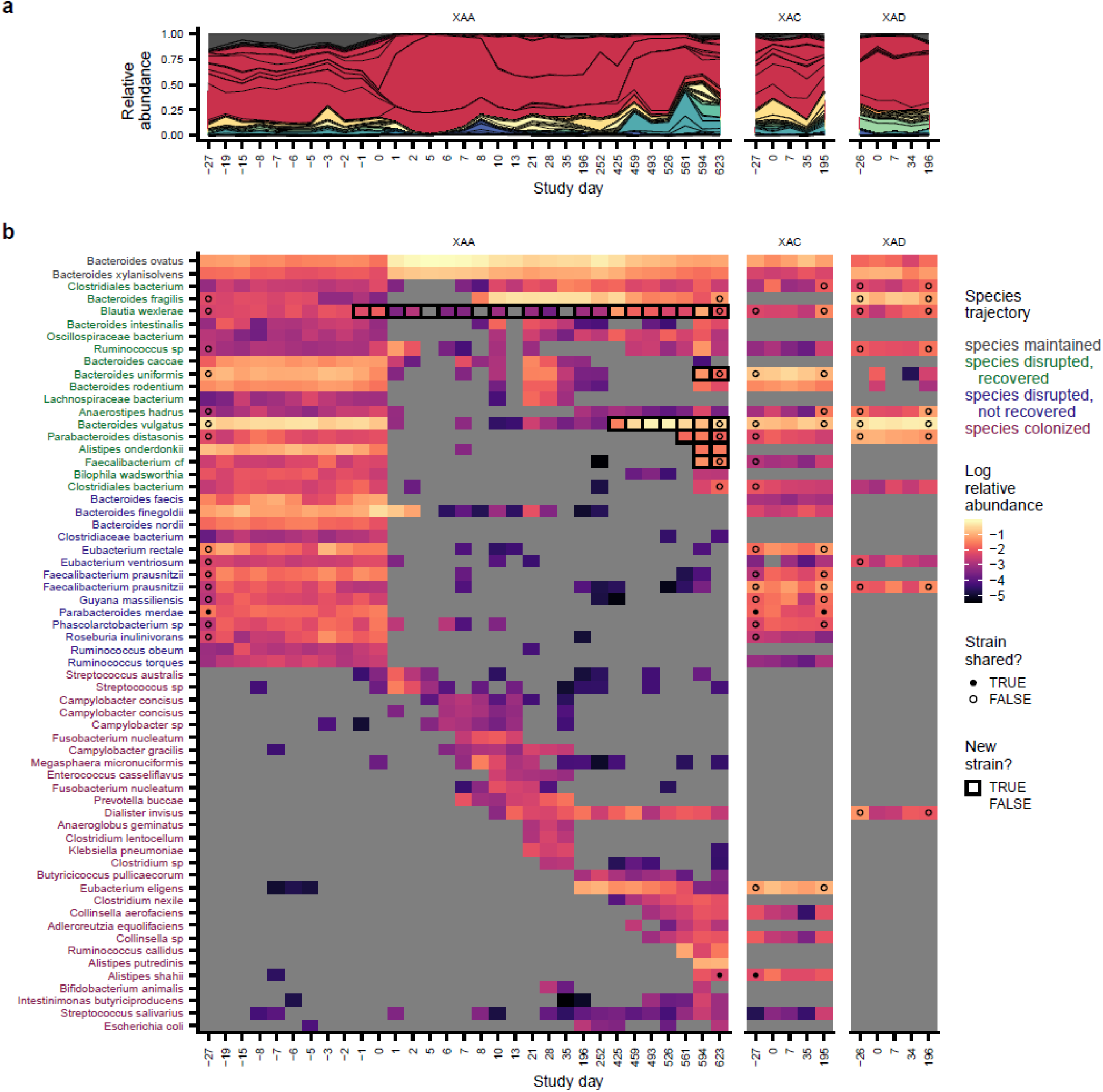
Species dynamics in subject XAA. **a**, Species-level community composition over time in subject XAA, a subject with lasting antibiotic responses, and two cohabiting controls, subjects XAC and XAD. **b**, Analogous version of **Fig. 2a** for subjects XAA, XAC, and XAD. Species are shown if they were present in XAA before antibiotics (median relative abundance >0.1%) or if they newly colonized XAA (**Methods**). Points at the first and last timepoints indicate whether the strains at that timepoint were shared (closed) or not shared (open) with any timepoint from the cohabiting subject (**Methods**); no point is shown if the species was not shared between subjects, or if the sequencing depth was insufficient to determine whether strains were shared. Timepoints are outlined in black if a new strain was detected relative to the initial timepoint.

**Extended Data Figure 5.**
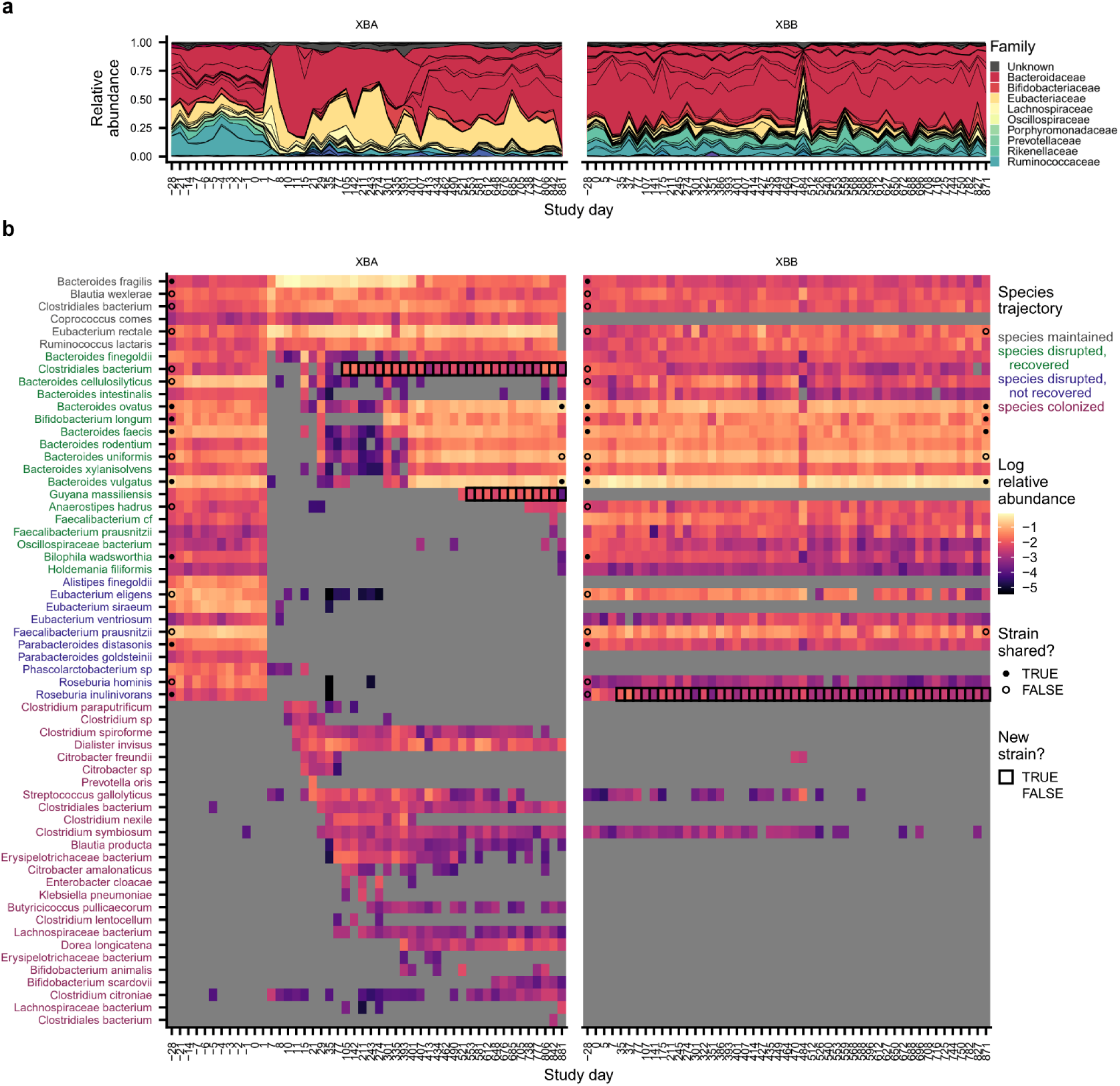
Species dynamics in subject XBA. Analogous version of **Extended Data** Fig. 4 for subject XBA. **a**, Species-level community composition over time in subject XBA, a subject with lasting antibiotic responses, and a cohabiting control, subject XBB. **b**, Relative abundances of species in subjects XBA and XBB. Species are shown if they were present in XBA before antibiotics (median relative abundance >0.1%) or if they newly colonized XBA (**Methods**). Points at the first and last timepoints indicate whether the strains at that timepoint were shared (closed) or not shared (open) with any timepoint from the cohabiting subject (**Methods**); no point is shown if the species was not shared between subjects, or if the sequencing depth was insufficient to determine whether strains were shared. Timepoints are outlined in black if a new strain was detected relative to the initial timepoint.

**Extended Data Figure 6.**
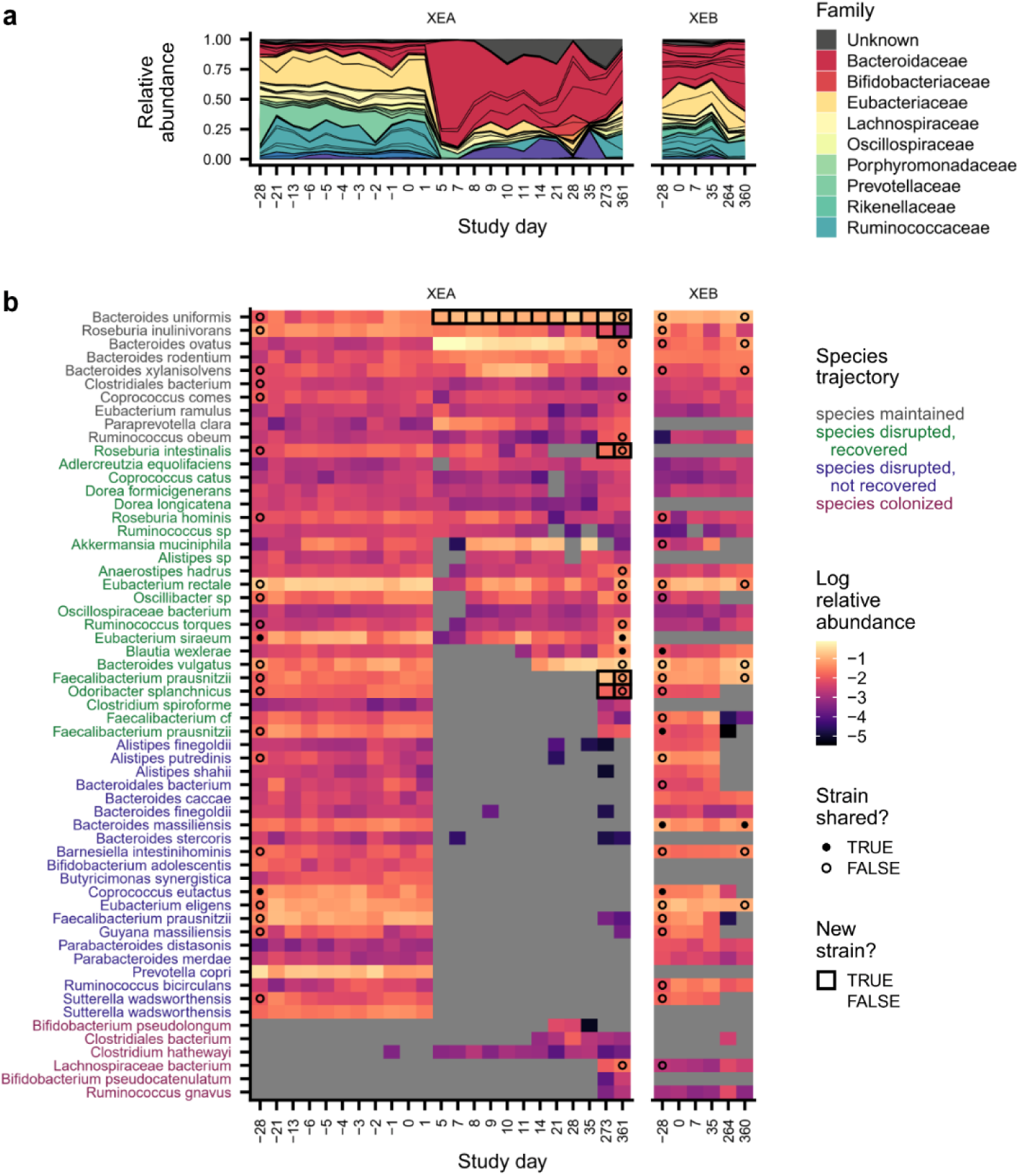
Species dynamics in subject XEA. Analogous version of **Extended Data** Fig. 4 for subject XEA. **a**, Species-level community composition over time in subject XEA, a subject with lasting antibiotic responses, and a cohabiting control, subject XEB. **b**, Relative abundances of species in subjects XEA and XEB. Species are shown if they were present in XEA before antibiotics (median relative abundance >0.1%) or if they newly colonized XEA (**Methods**). Points at the first and last timepoints indicate whether the strains at that timepoint were shared (closed) or not shared (open) with any timepoint from the cohabiting subject (**Methods**); no point is shown if the species was not shared between subjects, or if the sequencing depth was insufficient to determine whether strains were shared. Timepoints are outlined in black if a new strain was detected relative to the initial timepoint.

**Extended Data Figure 7.**
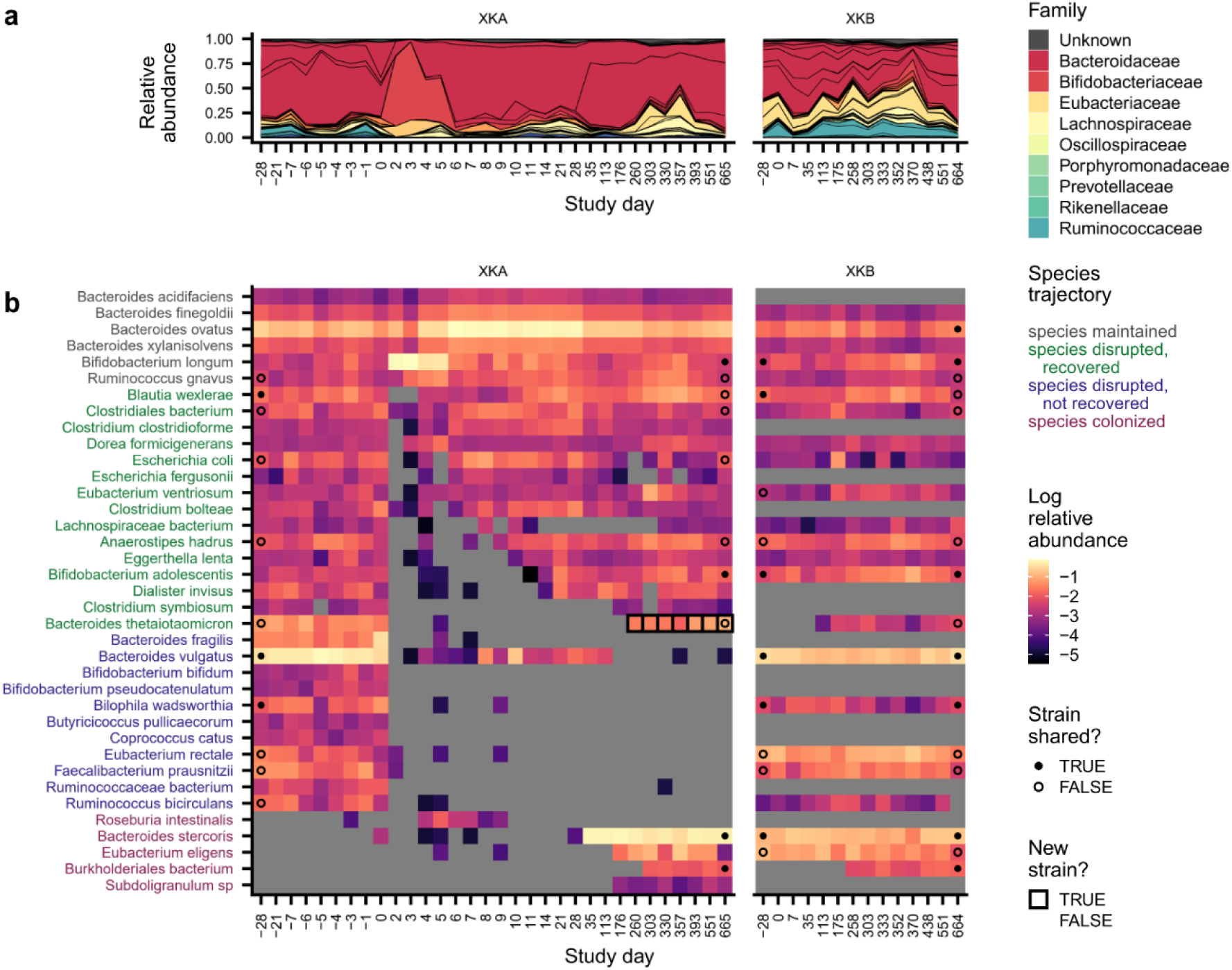
Species dynamics in subject XKA. Analogous version of **Extended Data** Fig. 4 for subject XKA. **a**, Species-level community composition over time in subject XKA, a subject with lasting antibiotic responses, and a cohabiting control, subject XKB. **b**, Relative abundances of species in subjects XKA and XKB. Species are shown if they were present in XKA before antibiotics (median relative abundance >0.1%) or if they newly colonized XKA (**Methods**). Points at the first and last timepoints indicate whether the strains at that timepoint were shared (closed) or not shared (open) with any timepoint from the cohabiting subject (**Methods**); no point is shown if the species was not shared between subjects, or if the sequencing depth was insufficient to determine whether strains were shared. Timepoints are outlined in black if a new strain was detected relative to the initial timepoint.

**Extended Data Figure 8.**
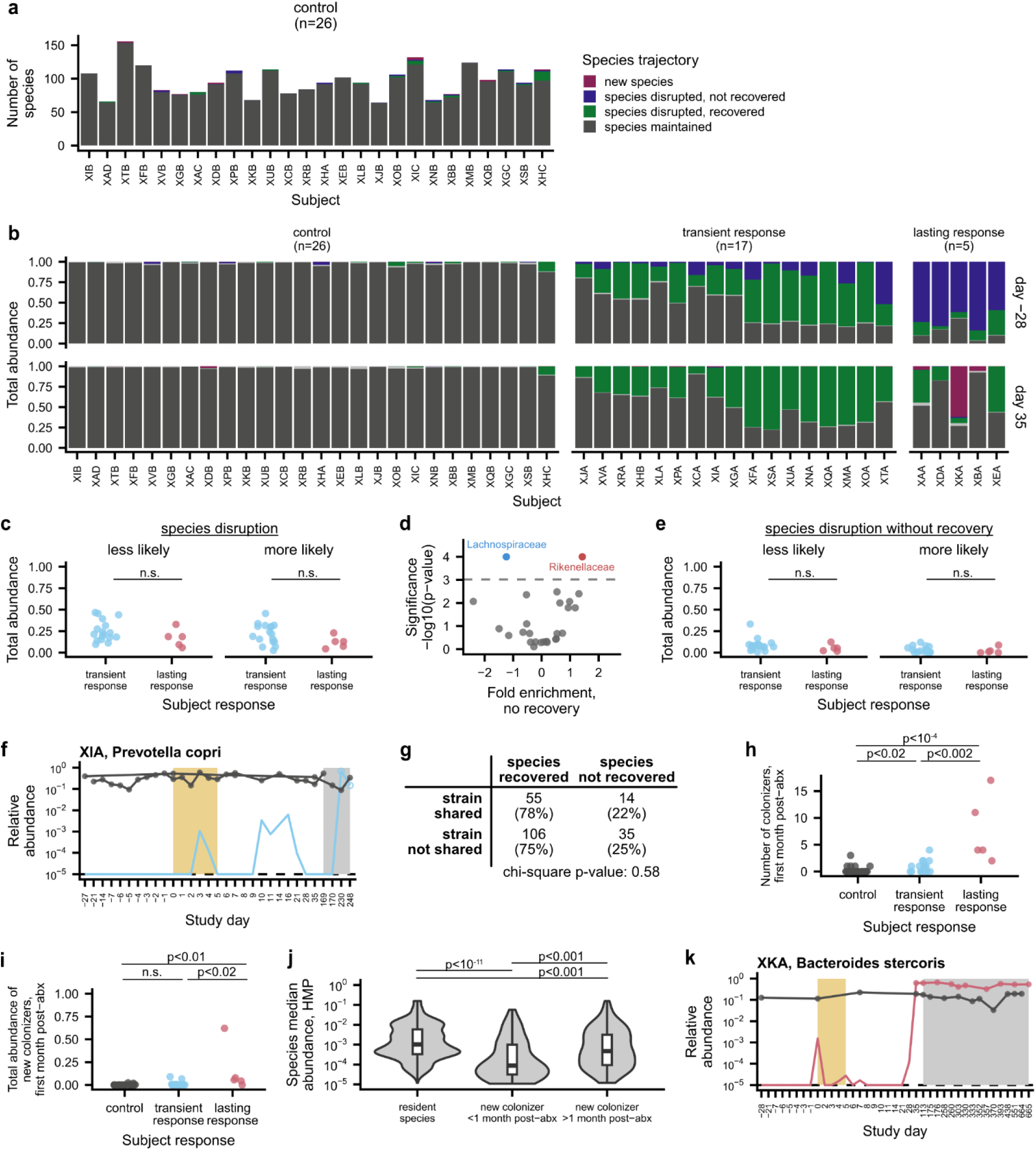
Species disruption, recovery, and colonization in the first month after ciprofloxacin perturbation. **a**, Number of species classified as maintained, disrupted, recovered, and colonized after ciprofloxacin in each control subject. **b**, Total abundance of species classified as maintained, disrupted, recovered, and colonized in each subject at the initial (day -28) and final (day 35) timepoints in the main study. **c**, The initial proportions of taxa that were significantly more or less likely to be disrupted were similar between subjects with transient and lasting antibiotic responses, suggesting that the ciprofloxacin sensitivity of individual taxa does not fully account for the heterogeneity across subjects. Points show the total abundance of bacterial families that were significantly less (left) or more (right) likely to be disrupted in each antibiotic-taking subject at the initial timepoint (Wilcoxon rank-sum two-sided tests, P>0.05). **d**, Enrichment and depletion of bacterial families among disrupted species that failed to recover. Families with significant enrichment (red) or depletion (blue) are labeled (**Methods**; permutation test, Bonferroni-corrected P<0.05). **e**, Total abundance of bacterial families that were significantly less (left) or more (right) likely to be disrupted without recovery in each antibiotic-taking subject at the initial timepoint (Wilcoxon rank-sum two-sided tests, P>0.05). **f**, Although colonization was rare in subjects with transient antibiotic responses, *Prevotella copri* colonized subject XIA (blue) and reached high relative abundances during follow-up sampling. Points indicate whether the strains at that timepoint were shared (closed) or not shared (open) with the strains at any timepoint in the cohabiting partner; points are not shown if the species was not shared between subjects, or if the sequencing depth was insufficient to determine whether strains were shared (**Methods**). The gold and gray boxes denote the ciprofloxacin perturbation and the period of follow-up sampling, respectively. The dashed line indicates the limit of detection of 10^-5^. **g**, Relationship between strain sharing and species recovery for subjects with transient antibiotic responses. Strain sharing between cohabiting subjects did not increase the likelihood of species recovery after ciprofloxacin perturbation (chi-square test for subjects with minimal and transient responses, n=210, χ^2^=0.3, P=0.58). **h**, Number of colonizing strains and species in each subject during the first month after ciprofloxacin perturbation (Wilcoxon rank-sum two-sided tests, controls vs. transient responders: n=43, r=0.36, P=0.02; controls vs. lasting responders: n=31, r=0.75, P=3.9 x 10^-5^; transient vs. lasting responders: n=22, r=0.67, P=0.002). **i**, Total relative abundance of colonizing strains and species in each subject during the first month after ciprofloxacin perturbation (Wilcoxon rank-sum two-sided tests, controls vs. transient responders: n=43, r=0.26, P=0.10; controls vs. lasting responders: n=31, r=0.68, P=1.9 x 10^-4^; transient vs. lasting responders: n=22, r=0.55, P=0.01). **j**, Median abundance of resident species and new colonizers in the Human Microbiome Project (Wilcoxon rank-sum two-sided tests, resident species vs. new colonizers <1 month post-antibiotics: n=1333, r=0.19, P=3.1 x 10^-12^; resident species vs. new colonizers >1 month post-antibiotics: n=1376, r=0.10, P=2.3 x 10^-4^; new colonizers <1 month post-antibiotics vs. new colonizers >1 month post-antibiotics: n=161, r=0.28, P=3.2 x 10^-4^). **k**, *Bacteroides stercoris* transmitted from subject XKB (gray) to subject XKA (red) after antibiotic perturbation.

**Extended Data Figure 9.**
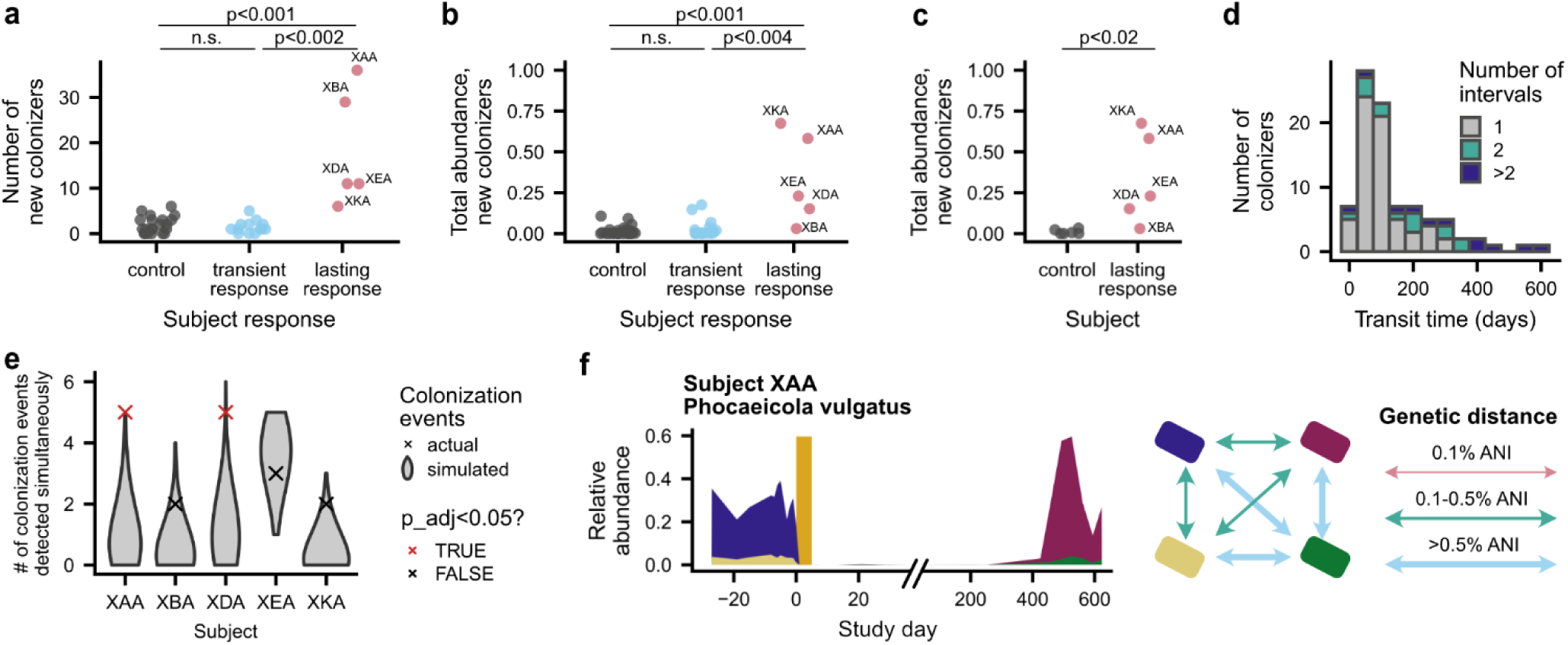
Colonization was elevated over longer timescales in subjects with lasting antibiotic responses. **a**, Number of strains and species that colonized each subject more than one month after the end of the ciprofloxacin course (Wilcoxon rank-sum two-sided tests, controls vs. transient responders: n=34, r=0.09, P=0.61; controls vs. lasting responders: n=26, r=0.67, P=7.0 x 10^-4^; transient vs. lasting responders: n=18, r=0.77, P=1.4 x 10^-3^). **b**, Total abundance of all new colonizers in each subject at the end of the study (Wilcoxon rank-sum two-sided tests, controls vs. transient responders: n=43, r=0.14, P=0.36; controls vs. lasting responders: n=31, r=0.60, P=9.5 x 10^-4^; transient vs. lasting responders: n=22, r=0.63, P=3.6 x 10^-3^). **c**, Total abundance of new colonizers in each subject with lasting antibiotic responses and their cohabiting partners at the end of the study (Wilcoxon rank-sum two-sided test, n=11, r=0.77, P=1.4 x 10^-2^). **d**, Distribution of transit times (the number of days between when a colonizing strain or species was last undetectable and when it reached steady-state relative abundance; **Methods**) across all new colonizers. Transit times were rounded up to the full time interval, regardless of the number of days between samples. Many species increased from undetectable to steady-state relative abundances within a single time interval and could be consistent with transit times as short as 1 day. **e**, Colonization events sometimes occurred in coordinated cohorts. Crosses show the maximum number of colonization events detected at a single follow-up sampling timepoint for each subject with lasting antibiotic responses. Colonization times were inferred by fitting relative abundance trajectories to a simple model of exponential growth (**Methods**). The gray distributions show the expected distribution if colonization were distributed uniformly over time (**Methods**; permutation test, subjects XAA and XDA, Bonferroni-corrected P<0.05). **f**, Two new strains of *Phocaeicola vulgatus* were detected simultaneously in subject XAA after antibiotic perturbation (**Methods**). Left panel shows the relative abundances of pre- and post-antibiotic strains (colored regions), while the right panel shows the genetic distance between them. The gold bar shows the timing of the ciprofloxacin perturbation.

**Extended Data Figure 10.**
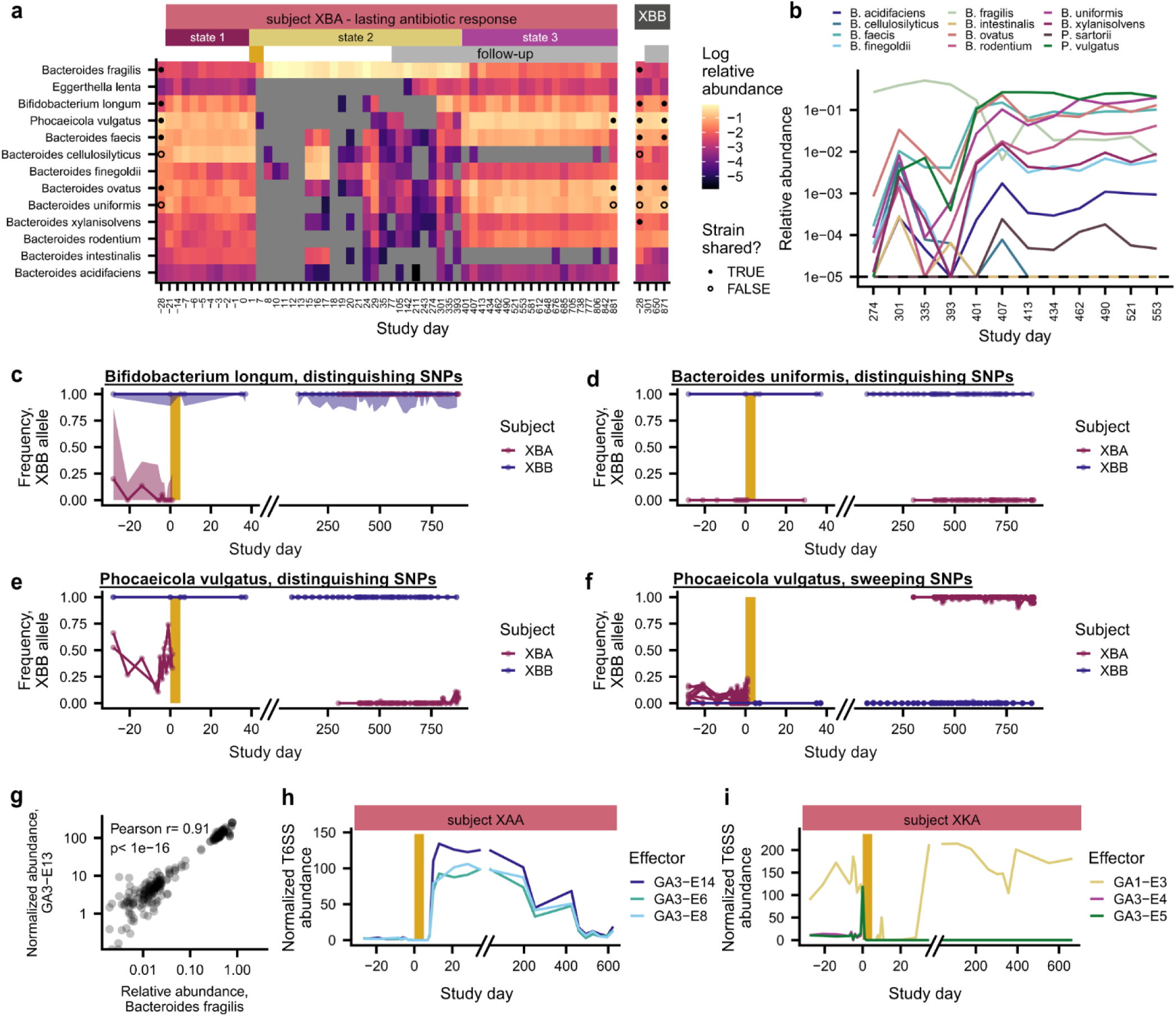
Strain and species dynamics in subject XBA. **a**, Subject XBA underwent rapid switches in community states. Relative abundances are shown for *Eggerthella lenta, Bifidobacterium longum,* and all *Bacteroidaceae* species detected in subject XBA. Strain sharing is annotated as in **Fig. 2** (**Methods**)**. b**, Several *Bacteroidaceae* species underwent sudden, simultaneous increases in relative abundance around day 400. **c,d**, Lines show the median abundance of SNPs that distinguish the initial strains of *Bifidobacterium longum* (**c**) and *Bacteroides uniformis* (**d**) in XBA and XBB. Shaded regions show the 25 and 75 percentiles of these SNP frequencies. **e**,**f**, Lines show the frequencies of the two SNPs that distinguish the initial strains of *Phocaeicola vulgatus* in XBA and XBB at the initial timepoint (**e**) and the seven SNPs that fixed in *Phocaeicola vulgatus* in XBA after the antibiotic perturbation (**f**). **g**, The relative abundance of *Bacteroides fragilis* was strongly correlated with the abundance of the T6SS GA3-E13 effector gene in subject XBA (Pearson’s test, n=210, ρ=0.91, P<2.2 x 10^-16^). **h,i**, Subjects XAA (**h**) and XKA (**i**) underwent major shifts in the abundance of T6SS genes after antibiotic perturbation.

## Notes

### Competing Interest Statement

The authors have declared no competing interest.

### Summary of Updates

Results section streamlined for ease of reading

